# The two-component regulator CvsR has a small core regulon *in planta* and modulates *Pseudomonas syringae* global gene expression with some overlap to the pattern triggered immunity stimulon response

**DOI:** 10.64898/2025.12.17.694939

**Authors:** Hsiao-Chun Chen, Carter J. Newton, Li Yang, Brian H. Kvitko

**Affiliations:** Department of Plant Pathology, University of Georgia, Athens, Georgia, USA; The Plant Center, University of Georgia, Athens, Georgia, USA

**Keywords:** Pattern-triggered immunity (PTI), *Pseudomonas syringae* pv. *tomato* DC3000, Two-component system, *in planta* transcriptome

## Abstract

Pattern-triggered immunity (PTI) provides broad-spectrum protection in plants by activating defense responses upon perception of conserved microbial signatures such as bacterial flagellin. *In vitro* transcriptome profiling revealed that the *Pseudomonas syringae* pv. *tomato* DC3000 two-component regulator CvsR mirrors some of the broader regulatory patterns observed under the exposure to PTI *in planta*. Our analyses indicated that during infection *in planta*, CvsR primarily governs a small core regulon centered on carbonic anhydrase and its associated transporter. Comparative RNA-seq analyses between the Δ*cvsR* and wild type strain further confirm this narrow regulatory scope. Moreover, the majority of bacterial transcriptional shifts appear to reflect indirect consequences of response to the host immune environment rather than direct CvsR-dependent regulation, including responses associated with sulfate starvation. Together, these findings suggest that PTI-driven bacterial transcriptional reprogramming is shaped predominantly by host immune status, with CvsR exerting modest, targeted control restricted to a limited set of genes.

## Introduction

Plants rely on a tiered innate immune system network to fend off pathogen attacks. Pattern-triggered immunity (PTI) and effector-triggered immunity (ETI) represent two functionally interdependent tiers of the plant immune systems (Jones and Dangl, 2006; Ngou et al., 2022). PTI is activated when cell-surface pattern recognition receptors (PRRs) directly bind to the conserved microbial molecules such as peptide epitopes of bacterial flagellin, elongation factor Tu, or cold shock proteins (Gómez-Gómez et al., 2001; Chinchilla et al., 2006; Kunze et al., 2004; Felix and Boller, 2003). Pathogens secrete specialized effector proteins that can suppress PTI or interfere with plant physiological responses. Resistant hosts harbor intracellular nucleotide-binding domain, leucine-rich-repeat receptors (NLRs) to recognize effectors directly or indirectly to trigger ETI. Core PTI signaling elements contribute to effective ETI, and ETI in turn enhances and sustains expression of PTI-associated genes (Yuan et al., 2021). PRRs are typically ligand specific form complexes with their co-receptors and trigger multiple signaling pathways including the mitogen-activated protein kinase (MAPK) signaling cascade, calcium dependent protein kinases pathways, and reactive oxygen species (ROS) (Ngou et al., 2022). Pre-activation of PTI by flg22 peptide 12 to 24 hours prior to bacterial infiltration results in potent immune outcome by limiting bacterial infections, effector delivery, and proliferation (Zipfel et al., 2004; Crabill et al., 2010). Upon activation of PTI, the apoplast undergoes rapid immune remodeling, including the accumulation of antimicrobial proteins and secondary metabolites (Chen et al., 2025; Lewis et al., 2015; Nobori et al., 2018; Zhou and Zhang, 2020; Miao et al., 2025). Additionally, PTI induces apoplast alkalization and alters ionic composition, further altering the physicochemical environment (Yu et al., 2019; Dora et al., 2022; Yang et al., 2024). These immune-driven changes are modeled to create a hostile microenvironment, requiring pathogens to deploy adaptive mechanisms for survival.

The bacterial pathogen *Pseudomonas syringae* pv. *tomato* DC3000 (*Pto* DC3000) is one of the most extensively studied model pathogens in plant–microbe interactions, largely due to its ability to infect the model plant *Arabidopsis thaliana* (Whalen et al., 1991). Virulence determinants of *Pto* DC3000, including the type III secretion system (T3SS), its repertoire of 36 effector proteins, and the phytotoxin coronatine (Buell et al., 2003; Xin et al., 2018), have been characterized in detail, establishing this strain as an ideal target for studying bacterial responses to PTI (Guo et al., 2009; Kvitko et al., 2009). RNA-seq, in particular, has overcome prior limitations by enabling deep, unbiased detection of both abundant and rare transcripts (Lee et al., 2017; Liao et al., 2019; Luneau et al., 2022). A critical methodological advance in this context using *Pto* DC3000 and *A. thaliana* pathosystem has been the physical enrichment of bacterial cells from plant tissue, which substantially increases the proportion of bacterial reads and achieves the mapping efficiency (Lovelace et al., 2018; Nobori et al., 2018; Wang et al., 2022).

While nutrient sequestration is widely proposed as part of PTI, direct evidence linking competition for these elements to bacterial growth inhibition remains limited (Herlihy et al., 2020; Dellagi et al., 2005; Rogan et al., 2024; Yamada and Mine, 2024; Fatima and Senthil-Kumar, 2021; Lovelace et al., 2018). These studies suggest that bacterial survival during immune activation may depend on the ability to sense and respond to host-induced environmental shifts, triggering adaptive transcriptional programs. Two component systems (TCSs) are widespread in Gram negative bacteria, functioning as core signaling modules that coordinate responses to environmental stimuli. A canonical TCS includes a membrane bound histidine kinase (HK) and a cytoplasmic response regulator (RR) that modulates gene expression upon phosphorylation. In *P. syringae*, the genomes encode over 60 TCSs signaling modules (Lavin et al., 2007). TCSs like PhoPQ, GacSA, RhpRS, CorRS, and CbrAB contribute to virulence and fitness *in planta* (Chatterjee et al., 2003; Deng et al., 2014; Fishman et al., 2018; Shao et al., 2021; Sreedharan et al., 2006). The CvsRS system has been linked to transcriptional responses also seen in response to pattern triggered immunity (PTI) (Fishman et al., 2018; Lovelace et al., 2018). Mutation of *cvsR* in *Pto* DC3000 show reduced colonization and virulence in tomato and *A. thaliana*, along with downregulation of type III secretion system genes, altered motility gene expression, and increased expression of sulfur uptake genes (Fishman et al., 2018). These changes mirror transcriptional trends in PTI exposed condition of *Pto* DC3000 (Lovelace et al., 2018). To better understand the contribution of CvsR to bacterial gene regulation during host immune activation, we analyzed the CvsR regulon with and without exposure to PTI. In this study, we perform RNA-seq on wild-type *Pto* DC3000 and Δ*cvsR* under PTI pre-activated and naïve apoplast conditions in *A. thaliana* to dissect the regulatory role of CvsR in bacterial response to PTI.

## Results and Discussion

### Wild type and *cvsR* mutant strains showed similar transcriptome profiles *in planta*

Transcriptomic studies of pathogens during early infection are inherently challenging due to the low proportion of pathogen-derived RNA relative to total host RNA. However, recent advances in tissue isolation methods, library preparation strategies, and sequencing technologies have markedly improved the ability to capture high-quality bacterial transcripts from infected plant tissues (Lovelace et al., 2018; Nobori et al., 2018; Honda et al., 2024). Physical separation of bacterial cells prior to RNA extraction produced a consistently high proportion of reads mapping to the bacterial genome (Lovelace et al., 2018; Wang et al., 2022). To investigate how CvsR modulates *Pto* DC3000 transcriptomic adaptation during PTI, 4.5-week-old *A. thaliana* Col-0 plants were pretreated with 1 μM flg22 to induce PTI or mock-treated with 0.1% DMSO as a naïve treatment for 16 h prior to syringe infiltration with either wild type or *cvsR* mutant strains. Bacterial RNA was isolated either directly from a centrifuged cell pellet of bacterial inoculum or from inoculated *A. thaliana* host tissue at 5 h post-inoculation (hpi) following a modified protocol based on Lovelace et al. (2018), which included vacuum infiltration with an RNA stabilization buffer followed by low-speed centrifugation to recover and enrich bacterial cells. Total RNA was extracted using a TRIzol-based method, subjected to both plant and bacterial rRNA depletion, and sequenced on the Illumina NovaSeq platform using 150 bp paired-end reads. Resulting reads were aligned simultaneously to the *A. thaliana* and *Pto* DC3000 reference genomes, enabling partitioning of host and bacterial transcripts and quantification of *in planta* bacterial gene expression. Using this approach, 40–97% of non-rRNA reads were maps to the *Pto* DC3000 genome across *in planta* libraries. On average, mock (naïve) “M” samples showed the higher mapping rate (93.5%), than flg22-pretreated PTI “F” samples (54%), while *in vitro* (pellet) “P” samples yielded 98.3% mapping (Figure 1<u>; Table S1</u>). Mapping rates to both bacterial and plant genomes were consistent across biological replicates. The removal of plant ribosomal RNA (rRNA) largely enhanced bacterial mRNA recovery was implemented in this study and resulted in substantially greater sequencing depth across *in planta* libraries than observed in Lovelace et al 2018.

**Figure 1.**
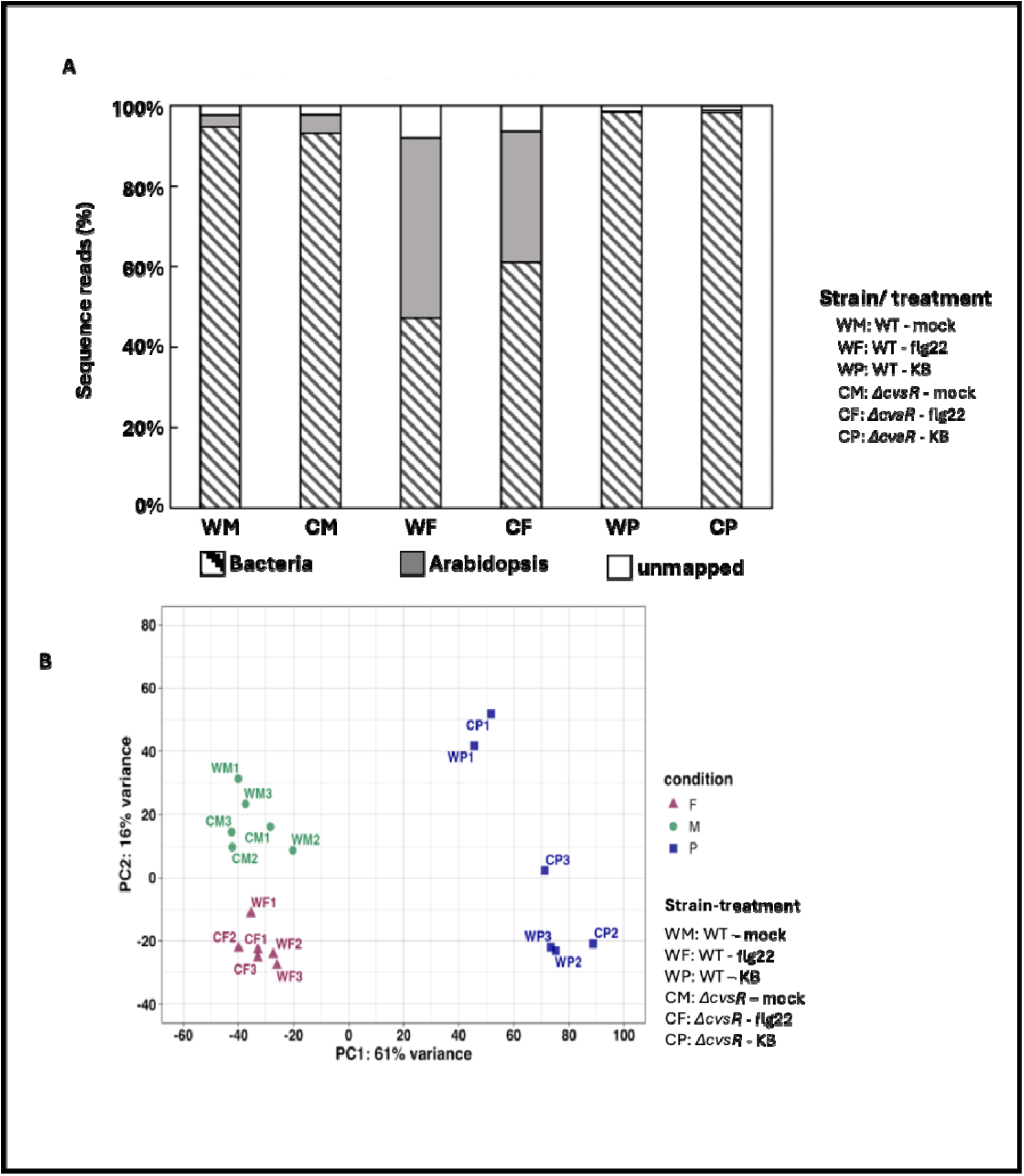
*Pseudomonas syringae* pv. *tomato* DC3000 (PtoDC3000)transcriptome profile in naïve and PTI condition of *Arabidopsis thaliana* leaves. (A) The proportions of sequencing reads mapped to *Pto* DC3000 genome, the *A. thaliana* genome, and unmapped reads are shown for all samples. Total RNA was extracted from leaves pretreated to induce pattern-triggered immunity (PTI) or from untreated naïve leaves, followed by inoculation with either the wild-type (WT) or Δ*cvsR* mutant strain of *Pto* DC3000 at 5 h post-inoculation (hpi) (n = 3). Additional RNA was isolated from bacterial starting inoculum from King’s B (KB) medium for both WT and Δ*cvsR* strains (n = 3). (B) *Pto* DC3000 transcriptome profile in *A. thaliana* at 5 hours post inoculation. Principal components analysis (PCA) of the gene-expression profile, measured in log-transformed bacterial gene counts (n = 3). Bacterial gene counts from sequenced total RNA samples inoculated pattern-triggered immunity–induced *A. thaliana* leaves (F: flg22) and inoculated naïve *A. thaliana* leaves (M: mock); *in vitro* bacterial culture from KB media (P)

Removal of plant and bacterial rRNA enabled robust detection of *Pto* DC3000 transcripts from inoculated leaf tissues with lower bacterial titer and sample quantities. This modification represents the highest mapping efficiency reported to date for *in planta* bacterial transcriptomic studies (Lovelace et al., 2018; Nobori et al., 2018; Honda et al., 2024) and holds potential promise for the transcriptome or genomic study of difficult patho-systems such as *Xylella* spp., Candidatus *Liberibacter* asiaticus (CLas), or Phytoplasma (De Francesco et al., 2022; Yang et al., 2025). However, physical separation of bacterial cells requires the use of structurally intact leaf tissue, as the apoplastic extraction procedure is particularly sensitive to mechanical disruption, especially during vacuum infiltration with the RNA-stabilizing buffer containing high salt concentrations. Consequently, this method may not be suitable for experimental contexts involving extensive tissue damage, such as during effector-triggered immunity (ETI) or late-stage disease progression. Together, these results demonstrate that optimized host rRNA depletion and careful bacterial enrichment enable reliable transcriptomic profiling of bacterial populations within plant tissues, even under immunity-activated states.

Principal component analysis (PCA) of normalized bacterial gene counts by DESeq2 analysis revealed that *in vitro* “P” samples for both wild type and Δ*cvsR* grown in KB medium were clearly distinct from *in planta* transcriptomes, suggesting a marked shift in bacterial gene expression between rich medium and the host environment (Figure 1B). Within the *in planta* samples, host immune status was the primary driver of transcriptional divergence: flg22-pretreated “F” samples clustered separately from mock (naïve) “M” samples, reflecting strong transcriptional reprogramming of bacteria in response to PTI. In contrast, wild type and *cvsR* mutant samples did not exhibit clear separation along the principal components and similar pattern was observed by sample distance clustering (Figure 1B; Figure S1), suggesting that host immunity exerts a stronger effect on bacterial transcriptional variation than the genetic factor of *cvsR* under these conditions. These global patterns (such as induction if motility genes) were consistent with previous *in planta* transcriptome studies from multiple laboratories, despite minor differences in duration of flg22 pre-treatment and bacterial inoculation (Lovelace et al., 2018; Nobori et al., 2018; Wang et al., 2022).

The lack of pronounced transcriptomic shifts in the *cvsR* mutant, even under *in vitro* conditions, contrasts with the regulatory model proposed by Fishman et al. (2018). However, variations in experimental conditions between the two studies may contribute to the difference. In the prior study, *cvsR* expression was induced by supplementation of nutrient broth with 5 mM calcium, whereas our analysis was performed under standard KB medium and *in planta* conditions. These differences in media composition, particularly the lack of supplemental calcium, may account for the observed differences. Notably, *cvsR* expression was comparable between *in planta* conditions and cultures grown in King’s B (KB) medium (Figure S2). In the leaf apoplast, *cvsR* mutation did not markedly alter the overall bacterial transcriptome despites the immune states of plant (Figure 2A). The expression level of *cvsR* remained insignificant between mock and PTI tissue (Figure S2). This implied that the host immune state remains the major determinant of bacterial gene expression during early stages of infection. Thus, *cvsR* and its regulon likely exert only a limited influence compared to the profound transcriptional reprogramming driven by host-derived cues in the early stage of plant colonization.

**Figure 2.**
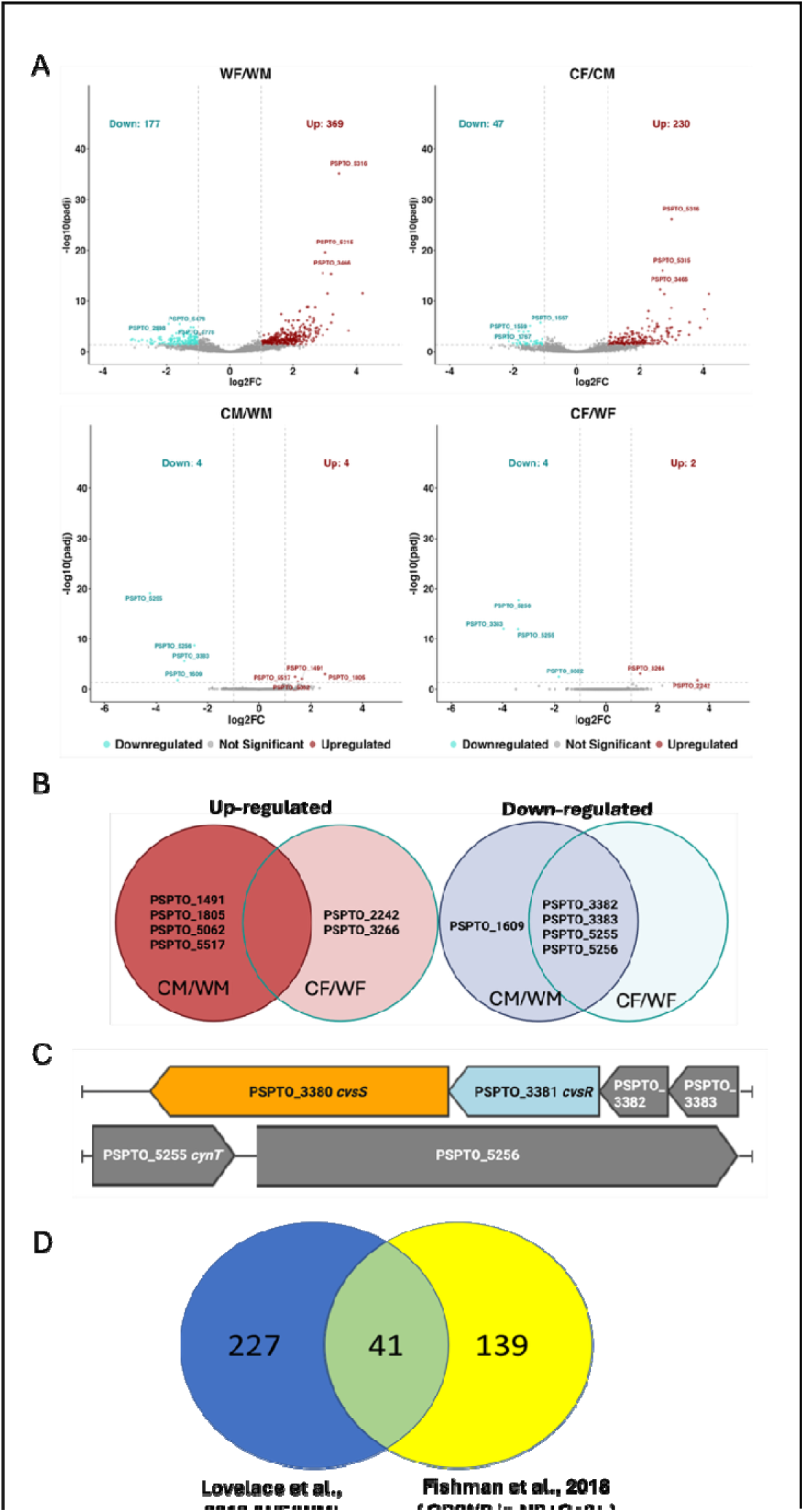
Comparison of *Pto* DC3000 transcriptomes from mock (M) or flg22 (F) treatments in wild-type (W) or Δ*cvsR* (C) strains. (A) Volcano plots display statistical significance (–log10 adjusted p-value) versus expression fold change (log2-transformed read counts). Red and blue points indicate significantly up- and down-regulated differentially expressed genes (DEGs). Top DEGs were highlighted. (B) Venn diagram showing comparisons of up- (red) and down (blue)-regulated DEGs between Δ*cvsR* and wild-type strains under both mock and flg22 treatments (C) Direct regulon of cvsR of two gene clusters (D) The overlap between transcriptomic DEGs from Lovelace et al. (2018) (blue; WF/WM, flg22 vs. mock at 5 hpi) and those from Fishman et al. (2018) (yellow; CP/WP, Δ*cvsR* vs. wild type under calcium-inducing *in vitro* conditions)

### The *cvsR* mutant has a reduced PTI stimulon response

To assess the role of CvsR in the bacterial PTI stimulon, we compared transcriptomic changes between wild type “W” and Δ*cvsR* “C” strains under mock “M”, flg22-pretreated “F”, and KB medium “P” conditions. In wild type *Pto* DC3000, the PTI stimulon (WF vs. WM) triggered extensive transcriptional reprogramming, with 546 differentially expressed genes (DEGs; |log2FC| > 1, padj < 0.05), including 369 upregulated and 177 downregulated genes (Figure 2A; <u>Table S2.1</u>). By contrast, the Δ*cvsR* mutant (CF vs. CM) exhibited an attenuated PTI stimulon response, with only 277 DEGs (230 upregulated, 47 downregulated) (Figure 2A; Table S2.2). The direct CvsR regulon, as previously determined based on the ChIP-seq analysis, revealed only very limited transcriptional changes with the activation of *cvsR* (Fishman et al., 2018). In naïve conditions (CM vs. WM), Δ*cvsR* displayed just 8 DEGs (4 upregulated, 4 downregulated) (Figure 2A; Table S2.3), whereas under PTI (CF vs. WF), only 6 DEGs were identified (2 upregulated, 4 downregulated) (Figure 2A; Table S2.4). Venn diagram comparisons highlighted distinct sets of *cvsR*-dependent genes under naïve “M” versus PTI “F” conditions. Specifically, four genes (PSPTO_1491, PSPTO_1805, PSPTO_5062, PSPTO_5517) were uniquely upregulated in the *cvsR* mutant under mock conditions, whereas two genes (PSPTO_2242, PSPTO_3266) were specifically upregulated under PTI conditions (Figure 2B). Conversely, three downregulated clusters (PSPTO_3382–3383, PSPTO_5255–5256, and PSPTO_1609) were shared between naïve and PTI conditions, suggesting stable CvsR-dependent repression across naïve and PTI tissue.

Among these, two directly regulated gene clusters—PSPTO_3382–3383 and PSPTO_5255–5256—were previously identified as direct CvsR targets via i*n vitro* ChIP-seq experiments (Fishman et al., 2018). Our data confirmed that these genes represent the core *cvsR* regulon. Gene schematic analysis showed that CvsS/CvsR directly regulate a putative operon containing the β-carbonic anhydrase (*cynT*; PSPTO_5255) and an adjacent major facilitator superfamily (MFS) transporter PSPTO_5256 (Figure 2C). Raw reads from RNA-seq indicated the depleted transcripts of both *cvsS*/*cvsR* and PSPTO_5255-5256 in Δ*cvsR* (Figure S2). Notably, the co-transcribed MFS transporter with the carbonic anhydrases was hypothesized to function in bicarbonate or carbonic acid transport to prevent surface-associated calcium phosphate precipitation and promote swarming ability (Fishman et al., 2019).

Despite the largely unchanged global transcriptome pattern between wild-type and Δ*cvsR* strains *in planta*, the reduced number of differentially expressed genes under PTI conditions (CF/CM) suggests that CvsR contributes to the bacterial PTI stimulon response. This finding indicates that while *cvsR* deletion does not broadly alter transcriptional profiles, CvsR still modulates specific transcriptional responses important for adaptation to the host immune environment. Direct CvsR-regulated genes remained responsive even *in planta*, verifying the *in planta* activity of this regulatory system and its limited overlap with other *cvsR*-associated genes reported previously (Fishman et al., 2018). These results also imply that *cvsR* and its direct regulon may play a more pronounced role in bacterial adaptation to environmental cues outside the plant apoplast.

*In planta*, the CvsS/CvsR system appears to primarily control a core operon comprising *cynT* and its adjacent transporter. This operon represents the only consistently regulated CvsR-dependent locus detected *in planta*, supporting its designation as the core *cvsR* regulon during host interaction. Given the hypothesized role of these proteins in bicarbonate or carbonic acid transport (Fishman et al., 2019), it would be informative to examine whether the global transcriptome profiles differ between *cvsR* and *cynT* mutants to further elucidate the functional significance of this regulatory module in bacterial adaptation to the host apoplast.

### CvsR-dependent modulation of PTI-associated pathways

To gain a systems-level view of the CvsR contribution to the PTI stimulon, we conducted KEGG pathway enrichment analyses across all experimental groups (Figure 3). The heatmap illustrates the strength of statistical confidence (–log q-value) for each pathway rather than absolute gene expression levels, thereby emphasizing the robustness of enrichment rather than the magnitude of transcriptional change or the number of genes in individual pathway. Comparison of Δ*cvsR* and wild-type strains under *in vitro* conditions (CP/WP) revealed only two significantly depleted pathways—bacterial chemotaxis and flagellar assembly—indicating reduced expression of motility-related genes in the absence of CvsR. Notably, CvsR-dependent regulation of motility-associated pathways was not observed *in planta*, under either naïve (CM/WM) or PTI (CF/WF) conditions, suggesting that the suppression of bacterial motility genes in planta override the regulatory effect by CvsR under *in vitro* condition. The second block of the heatmap highlighted KEGG pathways enriched in wild-type *Pto* DC3000 during PTI (WF/WM). Consistent with previous transcriptomic studies, PTI led to significant enrichment of metabolic and stress-response pathways, reflecting broad transcriptional reprogramming characteristic of the PTI stimulon. The third block summarized CvsR-dependent pathways under host-associated conditions. While the impact of CvsR was modest in naïve tissue (CM/WM), a stronger effect was observed during PTI (CF/WF), where pathway enrichment patterns were generally attenuated in the mutant relative to the wild type. This indicates that CvsR amplifies the magnitude and breadth of the PTI-responsive transcriptional network *in planta*.

**Figure 3.**
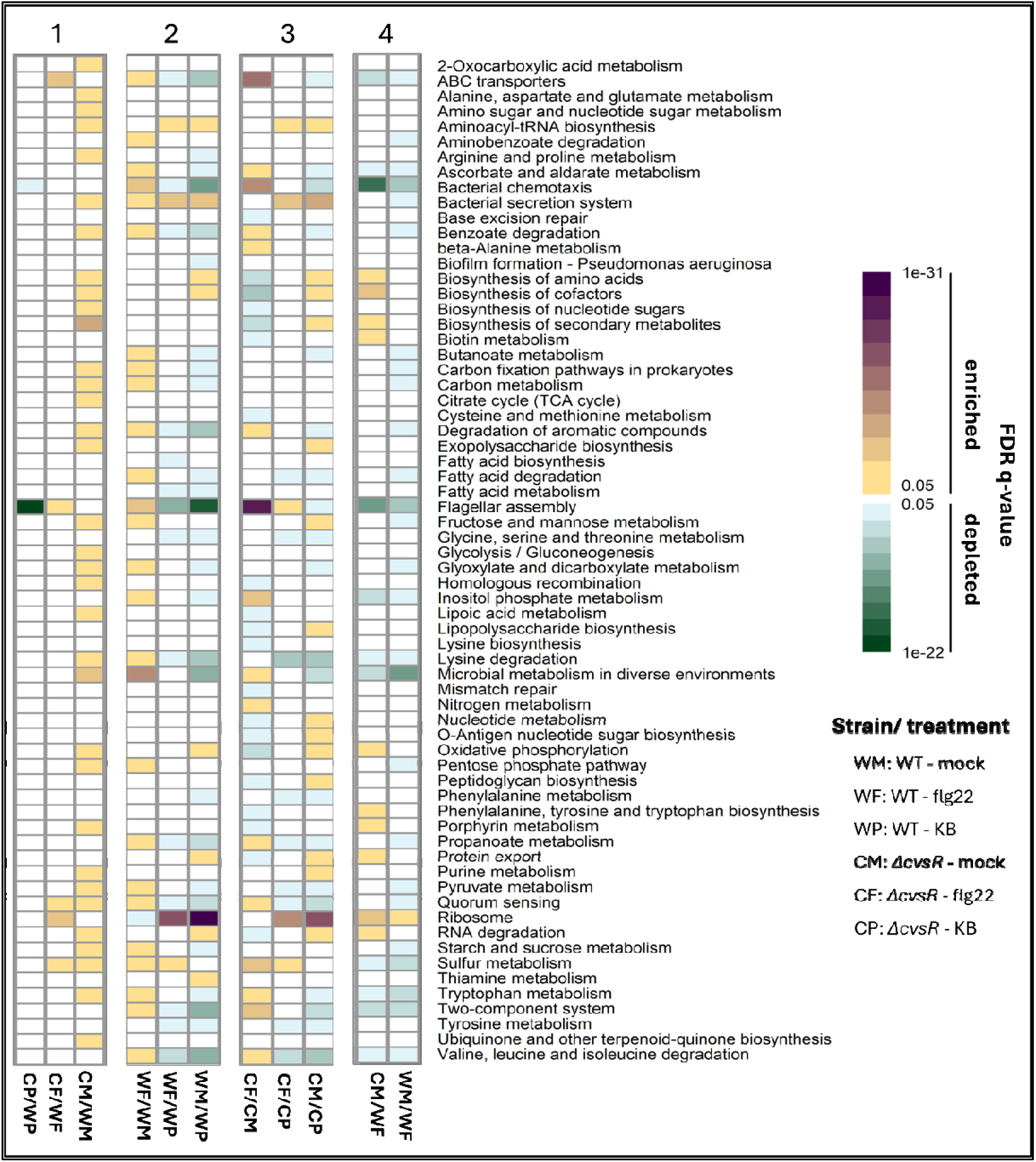
Differentially expressed KEGG (Kyoto Encyclopedia of Genes and Genomes) pathways for various comparisons in *Pto* DC3000. Heat map of q-values for KEGG pathways representing the differentially expressed genes in different comparisons of Δ*cvsR* mutant (C), wild type (W), flg22 treatment (F), and mock control (M). Differentially expressed pathways were identified using the goseq function in the gage v.2.24.0 R package. Color scale corresponds to the FDR-adjusted q-values, with darker shades indicating higher statistical significance. Pathways are classified as upregulated (purple scale) or downregulated (green scale) based on differential gene expression patterns.

The *in vitro* study by Fishman et al. (2018) identified under calcium induction in nutrient broth. The differentially expressed genes (DEGs) we observed (Lovelace et al., 2018) shared approximately 16% overlap with the PTI stimulon and 22% with the CvsR *in vitro* regulon (Figure 2D, Table S3). Based on this similarity, we hypothesized that the PTI may suppress CvsR signaling. Thus a *cvsR* mutant would have transcriptional responses mimicking those typically observed in response to the PTI stimulon. In the fourth block, we examined the Δ*cvsR* mutant in naïve tissue relative to the wild-type strain in PTI-activated tissue (CM/WF). If the Δ*cvsR* mutant showed PTI stimulon-like expression patterns under mock conditions, we would expect reduced transcriptional differences between Δ*cvsR* mock and wildtype flg22 conditions. However, a large number of significantly enriched pathways were still observed in the (CM/WF) comparison, and the expression pattern did not resemble that of the PTI regulon (WM/WF). This suggested that the regulation network of CvsR makes only limited contributions to the bacterial response to PTI.

### CvsR modulation of bacterial secretion systems

In our study, the type III secretion system (T3SS) genes are notably up-regulated during pattern-triggered immunity (PTI) as compared to naïve plant infections (Figure 4). This result diverges somewhat from the findings of Lovelace et al. (2018), where PTI induction prior to infection mildly suppress T3SS gene expression at 5h post-inoculation but is consistent with patterns observed in Smith et al 2018 where T3SS expression is still maintained higher level in PTI tissue than naïve tissue at later time points (24h post-inoculation).

**Figure 4.**
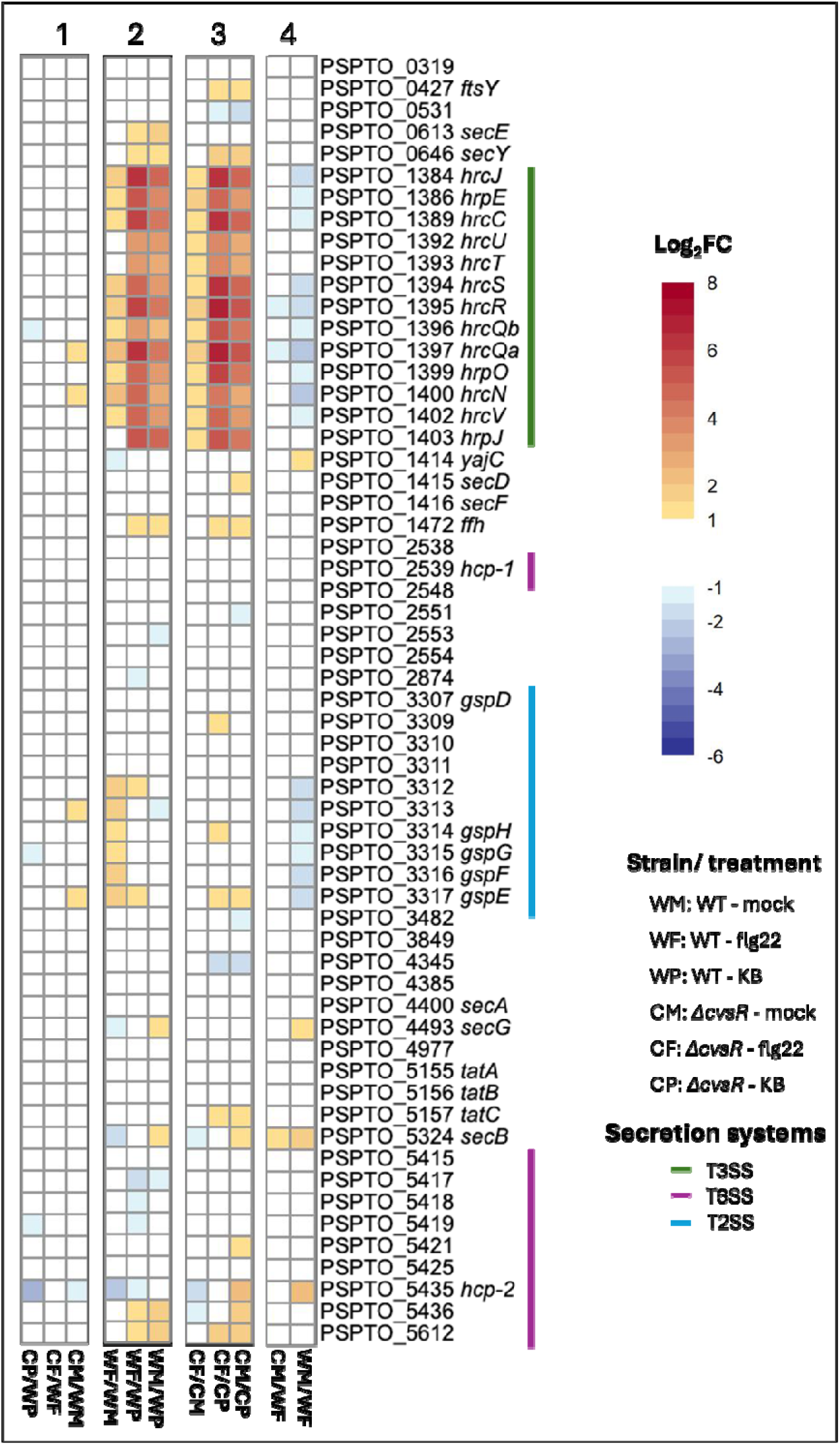
Gene expression level of bacterial secretion systems genes. Heat map displays the log2 fold change (log2FC) in mean normalized counts for *Pto* DC3000 bacterial secretion genes from the KEGG in different comparisons of *cvsR* mutant (C), wild type (W), flg22 treatment (F), and mock control (M). Colored cells indicate significant differential expression: red corresponds to upregulation and blue to downregulation (adjusted P-value < 0.05); white denotes genes not significantly changed (adjusted P-value > 0.05).

There was increased induction of T3SS genes in a *cvsR* mutant during PTI relative to the WT strain (CF/WF). *In vitro* expression of T3SS genes were lower in Δ*cvsR* than that in wildtype (Figure4, block1, CP/WP), aligning with the *in vitro* setting from previous report by Fishman et al., 2018. However, the reduction in T3SS genes in Δ*cvsR* was not observed *in planta* (Figure 4, block 1, CF/WF; CM/WM). The expression of type II secretion system (T2SS) pathway genes remained largely unchanged in wild-type strains during *in planta* colonization, but in the absence of *cvsR*, T2SS gene are upregulated (Figure 4, block 3).

Many plant-associated bacteria that possess the T6SS have strong competitive fitness when cultured with other bacteria, with few examples suggested T6SS contributed to the pathogen virulence (Wang et al., 2021; Kim et al., 2020). In the PTI tissue, most genes associated with T6SS, including *hcp-2*, were down-regulated in wildtype strain (Figure 4, block2, WF/WM, WF/WP). Mutation of *cvsR* diminishes the degree of T6SS downregulation and was even elevated in naïve condition compared to *in vitro* culture (Figure 4, block 3, CF/CP, CM/CP). Without *cvsR*, the PTI-induced suppression is lost, resulting in relatively sustained or less reduced levels of T6SS gene expression particularly in immune-challenged host tissue.

### CvsR modulation of bacterial sulfate and sulfonate importers and sulfur metabolism genes

The CvsS/CvsR two component system were previously observed to contribute to regulation of sulfate and sulfonate importers and sulfur metabolism (Fishman et al., 2018). While sulfur metabolism did not show significant changes at the pathway level in the *in vitro* comparison (Figure 3, CP/WP), transcriptome data revealed modest downregulation of individual sulfur metabolism–associated genes (e.g., PSPTO_0109, *ssuE*) in the Δ*cvsR* under *in vitro* conditions (Figure 5, block 1, CP/WP). However, this effect does not extend to the genes for sulfate and sulfonate periplasmic biding proteins *sbp* and *sfbp* (PSPTO_5316), contrasting with prior *in vitro* study by calcium augmentation (Fishman et al., 2018). *In planta*, this repression largely disappears, as indicated by negligible changes in log2 fold-change for gene expression across the entire sulfur metabolism pathway. The only notable exception is PSPTO_2590, which encodes a dimethylsulfone monooxygenase *sfnG*. Strong induction of genes involved in sulfate import including *sbp* and *sfbp* was detected during plant infection, with this upregulation being even higher in PTI-induced tissue, consistent with previous findings by Lovelace et al., 2018. The loss of *cvsR* further amplifies this response.

**Figure 5.**
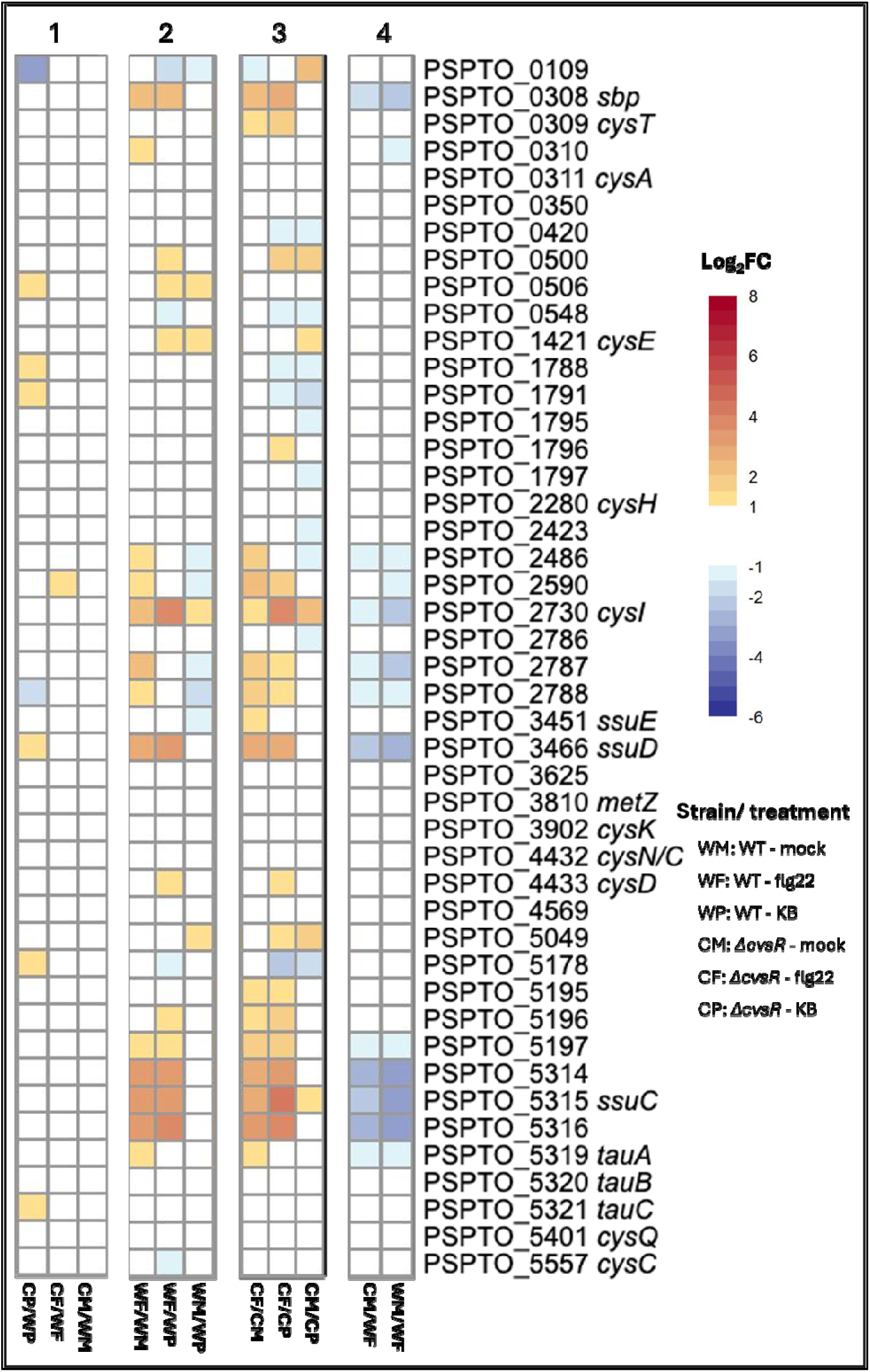
Gene expression level of sulfur metabolism genes. Heat map displays the log2 fold change (log2FC) in mean normalized counts for *Pto* DC3000 sulfur metabolism genes from the KEGG in different comparisons of *cvsR* mutant (C), wild type (W), flg22 treatment (F), and mock control (M). Colored cells indicate significant differential expression: red corresponds to upregulation and blue to downregulation (adjusted P-value < 0.05); white denotes genes not significantly changed (adjusted P-value > 0.05).

Sulfate metabolism genes are among the most distinct DEGs during bacterial responses to PTI (Lovelace et al., 2018). Our *in vitro* data showed no major reduction in the expression of the sulfate metabolism pathway (KEGG) in King’s B media (CP/WP), although several genes exhibited mild downregulation, consistent with the *in vitro* study by Fishman et al. (2018). However, this reduction was minor compared to the calcium-induced repression reported previously. Moreover, we did not observe strong suppression of *sbp* and *sfbp*, which were the least affected sulfate metabolism genes in the *cvsR* mutant. Sulfate metabolism genes exhibited higher expression under PTI stimulation (Figure 5, WF/WM, WF/WP, CF/CM, CF/CP; Figure S3), which likely explains the stronger induction observed in the *cvsR* mutant *in planta* compared to *in vitro*, reflecting *cvsR*-dependent downregulation of this pathway under *in vitro* conditions.

### CvsR modulation of bacterial chemotaxis and flagellar regulation

Distinctive patterns of bacterial chemotaxis and flagella assembly genes under *in vitro* and *in planta* conditions provided evidence of limited impact of *cvsR* during plant infection (Figure 6). Under *In vitro* condition, chemotaxis and flagella pathways were downregulated in the Δ*cvsR* compared to the wild type (Figure 6, block1, CP/WP), consistent with the impaired motility observed in Δ*cvsR.*This finding corroborates previous reports demonstrating a cvsR-dependent swarming phenotype and defect in motility upon *cvsR* disruption (Fishman et al., 2018; Fishman et al., 2019). However, when these genes are examined *in planta*, regardless of immune status (Figure 6, block 1, CM/WM and CF/WF), chemotaxis and motility gene expression does not significantly differ between Δ*cvsR* and wild type. Surprisingly, while flagella and chemotaxis genes are much less expressed *in planta* than *in vitro* in the wild type strains (Figure 6, block 2), the expression of many genes (e.g*. motA-2*, *mot-B*, *fliC*) displayed up-regulation *in planta* than *in vitro* with the mutation of *cvsR* with higher expression level in PTI compared to naïve tissue (Figure 6, block 3).

**Figure 6.**
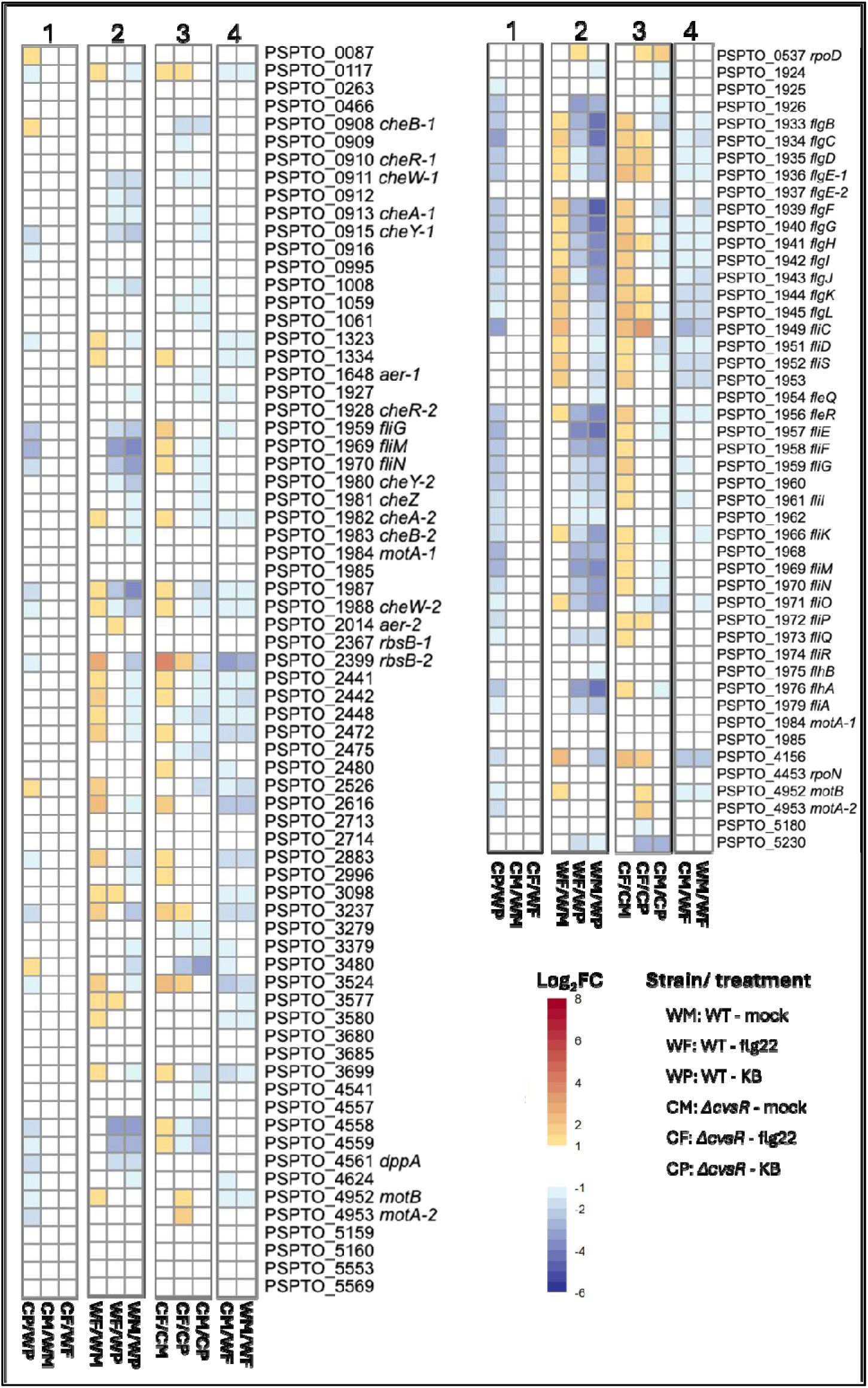
Gene expression level of motility genes. Heat map displays the log2 fold change (log2FC) in mean normalized counts for *Pto* DC3000 from the KEGG in different comparisons of *cvsR* mutant (C), wild type (W), flg22 treatment (F), and mock control (M). Colored cells indicate significant differential expression: red corresponds to upregulation and blue to downregulation (adjusted P-value < 0.05); white denotes genes not significantly changed (adjusted P-value > 0.05).

In contrast to the inducing conditions under which 199 genes were reported as potential direct targets of CvsR (Fishman et al., 2018), our data suggest that CvsR exhibits a highly restricted core regulon *in planta*. This regulon appears to be confined to autoregulation of the *cvsS*/*cvsR* two-component system and regulation of *cynT* (carbonic anhydrase) and its associated transporter PSPTO_5256, with no evidence for direct regulation of additional transcription factors. Thus, the subtle CvsR modulation of multiple pathways including motility and sulfur metabolism may be the results of altered expression of carbonic anhydrase expression. Carbonic anhydrase may reduce local pH on the surface of cells through production of carbonic acid (Fishman et al., 2019). Thus, the reduced *cynT* expression in Δ*cvsR* may result in increased local pH around colonization zone in apoplast. During PTI, the apoplast undergoes a significant pH increase (Yang et al., 2024). Thus, the observed partial overlap between a *cvsR* modulated genes and the response to the PTI stimulon may be attributable to bacterial responses to increased local pH.

## Conclusion

In this study, we established an optimized *in planta* bacterial transcriptome profiling approach that achieves exceptionally high bacterial read recovery through combined host rRNA depletion and physical enrichment. This method enables reliable resolution of bacterial transcriptional dynamics under immunity-activated states. Our results demonstrate that while CvsR contributes minimally to global transcriptional reprogramming within the host, it retains distinct regulatory activity over a small, conserved regulon, primarily encompassing the β-carbonic anhydrase (PSPTO_5255) and its adjacent MFS transporter (PSPTO_5256) *in planta*. Under *in vitro* conditions, CvsR regulates motility- and sulfur metabolism–associated pathways, consistent with previous reports, but these effects are largely attenuated *in planta*. The attenuated PTI stimulon responses observed in the Δ*cvsR* mutant further suggests that CvsR fine-tunes bacterial responses to immune-induced environmental cues rather than acting as a major transcriptional driver. Collectively, these findings indicate that CvsR functions as a context-dependent regulator with limited influence in the plant apoplast, yet its role may be more pronounced in mediating bacterial adaptation to external or pre-infection environments.

## Material and methods

### Plant tissue preparation and growth condition

*Arabidopsis thaliana* (Col-0) seeds were sown under the mesh-covered pot as described previously (Lovelace et al., 2018). Plants were grown in a growth chamber (Conviron A1000) for 4.5 weeks under long-day conditions at 23°C (14-h day and 10-h night) at 70 μmol light settings. One day prior to treatment, plants were removed from the growth chamber and were placed in a growth room under 12-h day and 12-h night conditions. 3 to 5 fully expanded leaves on each plant on each plant were infiltrated with either 1 μM flg22 in 0.1% DMSO or with 0.1% DMSO by 1-ml blunt syringes on the abaxial surface. Plants were kept in the same growth room for 16 h prior to bacterial inoculation.

### Bacterial inoculation

*Pseudomonas syringae* pv. *tomato* DC3000 (*Pto* DC3000) was prepared as lawn, growing on King’s B (KB) (King et al. 1954) agar supplemented with 40 μg/mL rifampicin and were grown overnight at 28°C. Inoculum was harvested from plates by washing out bacterial smear from the plate surface and suspended and washed twice in 0.25 mM MgCl_2_ .OD_600_ of 0.8 (1 × 10^9^ CFU/mL) was reached as the final concentration for inoculum. Bacterial inoculum was infiltrated with a blunt-end syringe into treated leaves as described above. Plants were allowed to dry at ambient temperature (22°C) on bench for 5 hours before sampling.

### Extraction of bacteria from infected plants

At 5 hpi, leaves from each treatment were harvested by cutting at the leaf blade junction. Isolation of bacteria from leaf apoplast was followed from the previous method (Lovelace et al., 2018) with modification. Briefly, 80-100 leaves were arranged on four sheets of parafilm, rolled and inserted into the barrels of four 20-ml syringes. An ice-cold RNA stabilizing buffer (Invitrogen) was poured into each syringe, which was sealed and vacuum infiltrated at 95 kPa for 2 min, followed by a slow release of the vacuum. The RNA stabilizing buffer was spun out from apoplast at 1000 × g for 10 min at 4°C to isolate bacterial RNA.

The flow-through was pooled for each biological replicate form each treatment and was concentrated by syringe filtration, using a 0.20-μm Micropore membrane (Millipore). Filters were placed in homogenization tubes, were flash frozen in liquid nitrogen, and were stored at −80°C for RNA extraction.

### Bacterial RNA isolation and sequencing

The filter membranes were homogenized in Trizol reagent (Thermo Fisher Scientific, Waltham, MA) for 1 min at 1,750 Hz, using a Geno/Grinder (Thermo Fisher Scientific, Waltham, MA) with three x 3 mm high density zirconium beads followed by the chloroform-ethanol isolation from manufacture (TRIzol™ Reagent User Guide, Pub. No. MAN0001271 D). For *in vitro* samples, 1 ml of the 1 × 10^9^ CFU/ml bacterial inoculum in 0.25 mM MgCl_2_ was pelleted and RNA was extracted followed by the same procedure above.

RNA samples were additionally treated with TURBO DNase (Invitrogen, Carlsbad, CA) to eliminate genomic DNA contamination. RNA was quantified using the NanoDrop OneC Microvolume Spectrophotometer (Thermo Fisher Scientific, Waltham, MA). Three independent bacterial suspensions started from three colonies were sampled for a total of three biological replicates. The plant samples were depleted of rRNA using the QIAseq FastSelect –rRNA Plant Kit (Qiagen, Germantown, MD) and bacteria samples were depleted of rRNA using the QIAGEN FastSelect rRNA HMR Kit (Qiagen, Germantown, MD). Next, RNA sequencing libraries were constructed with the NEBNext Ultra II RNA Library Preparation Kit for Illumina by following the manufacturer’s recommendations (New England Biolabs, lpswich, MA). Paired-end 150-nt reads were sequenced using the Nova-seq platform (Illumina, San Diego, CA).

### RNA-Seq data analysis

Reads were quality trimmed using Trimmomatic (Bolger et al., 2014) and were aligned to the RefSeq *P. syringae* pv. *tomato* DC3000 genome (NC_004278.1, NC_004632.1, NC_004633.1), using Bowtie2 (Langmead and Salzberg et al., 2012). Counts of RNA-Seq fragments were computed for each annotated gene, from reads per kilobase million values, using the stringtie script (Pertea et al., 2016). DEGs were identified from gene counts for each sample, using the Bioconductor package DESeq2 version 3.21 (Love et al., 2014). DEGs were selected based on log2-transformed and normalized mean counts that have an adjusted P value below a false discovery rate (FDR) cutoff of 0.05. DEGs set from each comparison were extracted with additional cutoff of |log2FC|>1.0.

Principal components analysis of log2-transformed normalized counts was performed for all treatments and replicates, using the rlog function in DESeq2. Volcano plot was performed on log2 fold changes in mean normalized counts between treatments using DESeq2, and heatmaps were generated using the R package pheatmap version 1.0.13.

KEGG analysis was conducted using log2 fold change values of normalized mean counts for annotated genes using the Bioconductor package, gauge version 2.24.0 (Luo et al., 2009). Gene sets of metabolic pathways were obtained, from the KEGG pathway database, using the organismal code “pst” for *Pto* DC3000. Significant gene sets were identified from log2 fold changes between treatments at similar timepoints and were selected based on a FDR q value cut-off of 0.05. The data represented enrichment of pathways rather than the expression level between the treatment pair.

### qPCR analysis on selected genes of interest

Four replicates of 4.5-week-old *A. thaliana* Col-0 plants were treated with 1 μM flg22 in 0.1% DMSO or 0.1% DMSO 16 h prior to inoculation as the same manner described above. The bacterial inoculum of *Pto* DC3000 wild type or *cvsR* mutant was prepared as described above and was further diluted to a final concentration of approximately 1 × 10^9^ CFU/ml and was infiltrated into marked leaves of all plants. At 5 hpi, inoculated leaves from a single plant were harvested and homogenized. Ground tissues were set for RNA extraction using TriZol and TURBO-DNase cleanup described as above. Four independent samples of pelleted initial inoculum were also flash-frozen in liquid nitrogen for RT-qPCR analysis. cDNA synthesis, RT-qPCR, and normalization were conducted based on previously described procedures (Smith et al. 2018).

Normalized cDNA was tested from four treatments, i.e., wild type/*cvsR* mutant strain from KB, naïve host tissue, and PTI tissue. These samples were tested for relative expression of two genes of interest *sbp*, and *sfbp*. Samples were grouped by biological replicate set and were tested against five genes of interest, two previously validated *recA* as reference gene, inoculum, and *Pto* DC3000-specific 16S rRNA within the same plate (Smith et al. 2018). Relative expression of in-planta samples normalized to the inoculum sample measured as the NRQ of a gene of interest was calculated as described previously (Smith et al. 2018). Relative expression of bacteria exposed to PTI relative to bacteria during naïve host infection measured as the NRQ of a gene of interest was calculated similarly to that above; A student t-test was performed on log2 NRQ values for each gene of interest against wild type sample in naïve tissue (WM).

## Data Statement

The RNA-seq data used in this study are deposited in the National Center for Biotechnology Information Gene Expression Omnibus database.

## Acknowledgements

This work was supported by the National Science Foundation Grant 1844861 to B.H.K. L.Y. and C.J.N. were supported by the National Institutes of Health (R35GM143067 to L.Y.). We thank Dr. Melanie Filiatrault for providing *Pto* DC3000 Δ*cvsR* strain, and providing helpful feedback and comments.

## Author contributions

H-C.C. and B.H.K. designed the research; H-C.C. conducted experiments; H-C.C. and C.J.N. analyzed the data; H-C.C. and B.H.K. wrote the manuscript. All listed authors reviewed and approved draft and final versions of the manuscript.

## Supporting Materials

**Table S1.1 Sequences mapping rate to *Pseudomonas syringae* pv *tomato* DC3000 genome**

**Table S1.2 Sequences mapping rate to *Arabidopsis thaliana* col-0 genome**

**Table S2.1 significant DEGs between flg22 and mock treatment in wild-type strains (WF/WM)**

**Table S2.2 significant DEGs between flg22 and mock treatment in Δ*cvsR* strains (CF/CM)**

**Table S2.3 significant DEGs of Δ*cvsR* and wild-type strains under mock treatments (CM/WM)**

**Table S2.4 significant DEGs of Δ*cvsR* and wild-type strains under flg22 treatments (CF/WF)**

**Table S3. Concordantly regulated overlapping DEGs between the Lovelace and Fishman datasets**

**Figure S1.**
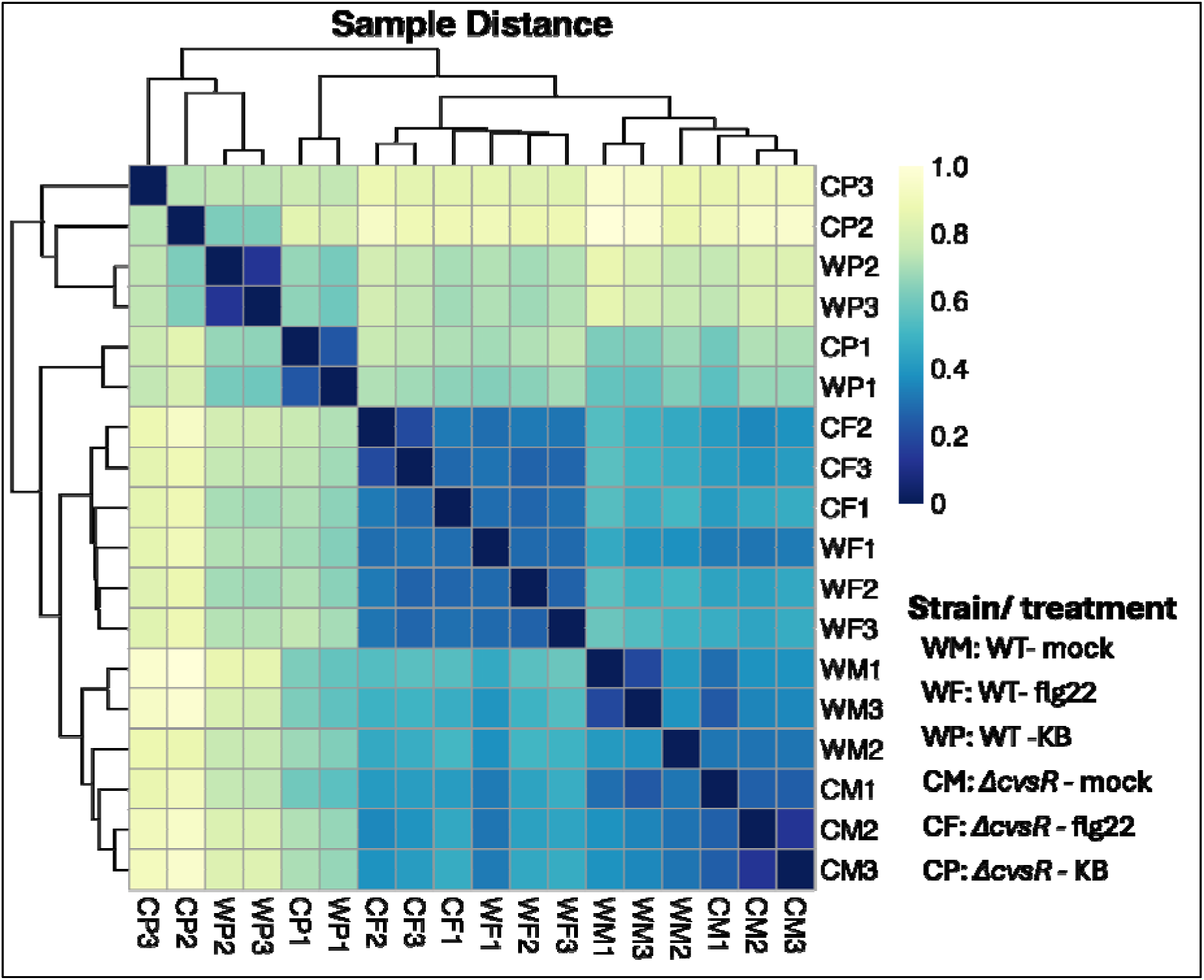
Expression heatmap representing sample-to-sample distances calculated from the variance-stabilizing transformation of RNA-seq count data across all genes. Hierarchical clustering of 18 RNA-seq samples provides an overview of relationships among genotypes, with sample proximity indicated by the intensity of square colors. Blue areas correspond to closely related samples with similar expression profiles, while yellow regions highlight greater sample-to-sample dissimilarity.

**Figure S2.**
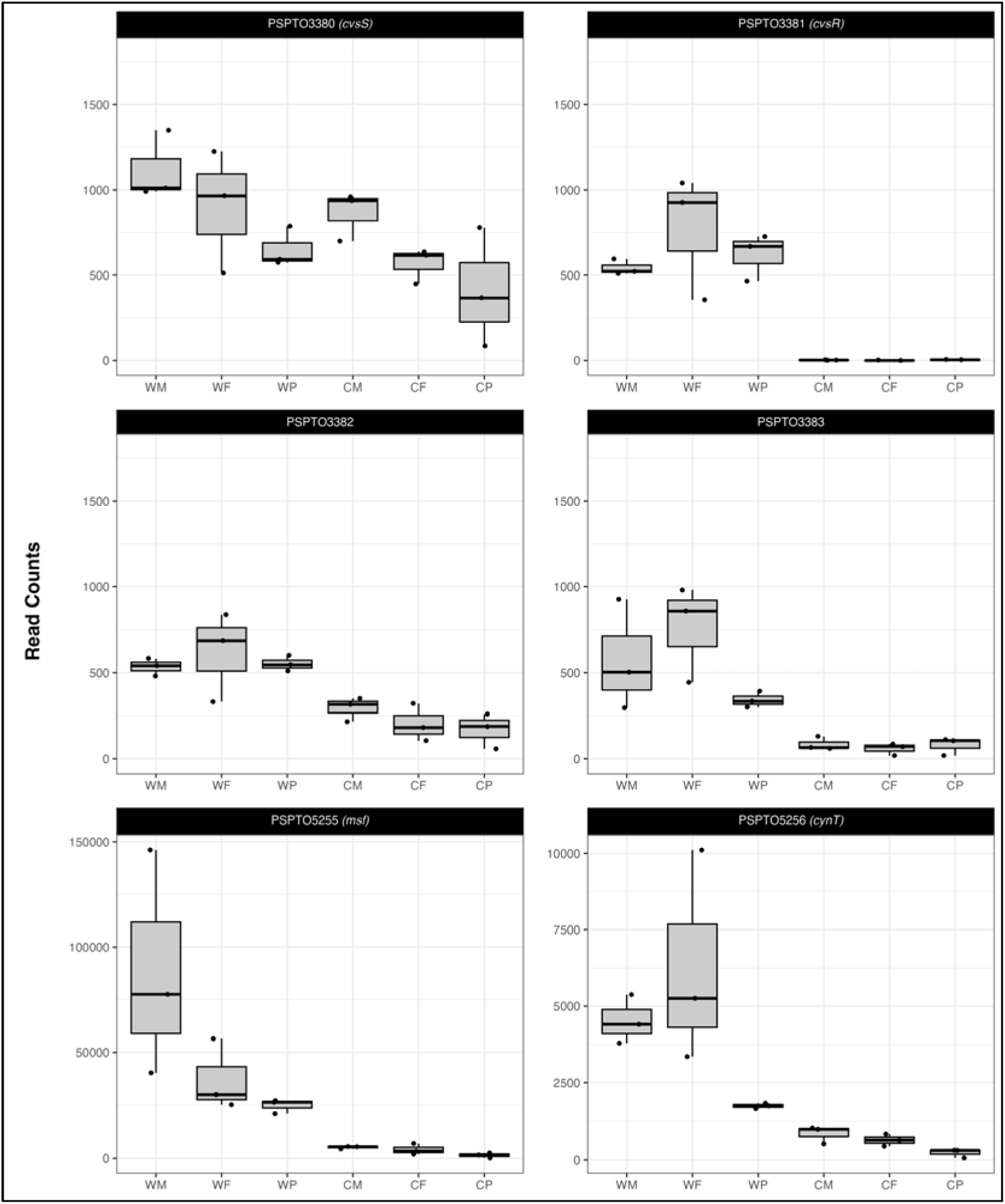
Raw read counts per sample of CvsR direct regulon. The y-axis represents the read counts and the x-axis represents sample type (strain/treatment). Each boxplot shows the distribution of read counts for three biological replicates (n=3) within each sample type.

**Figure S3.**
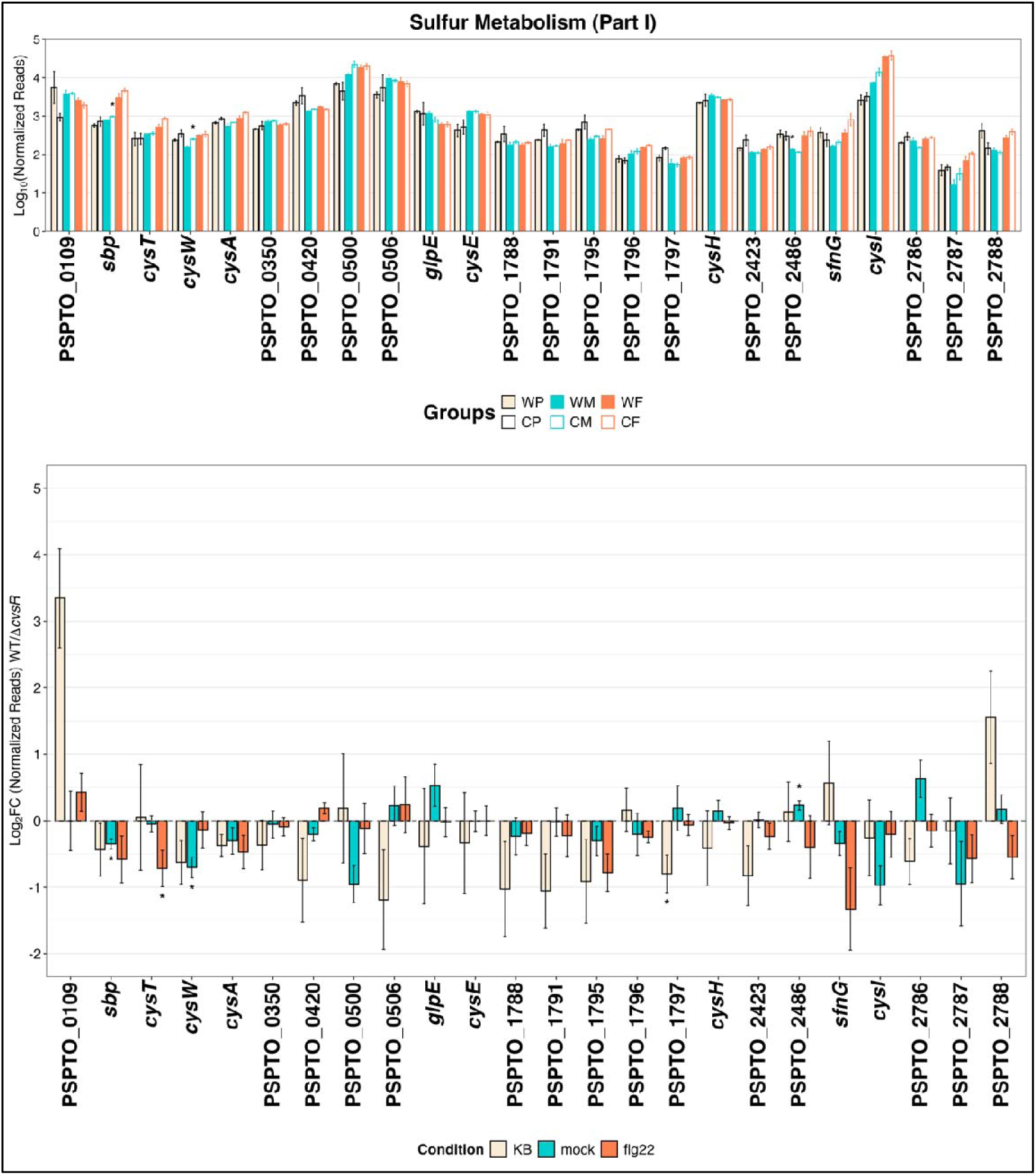
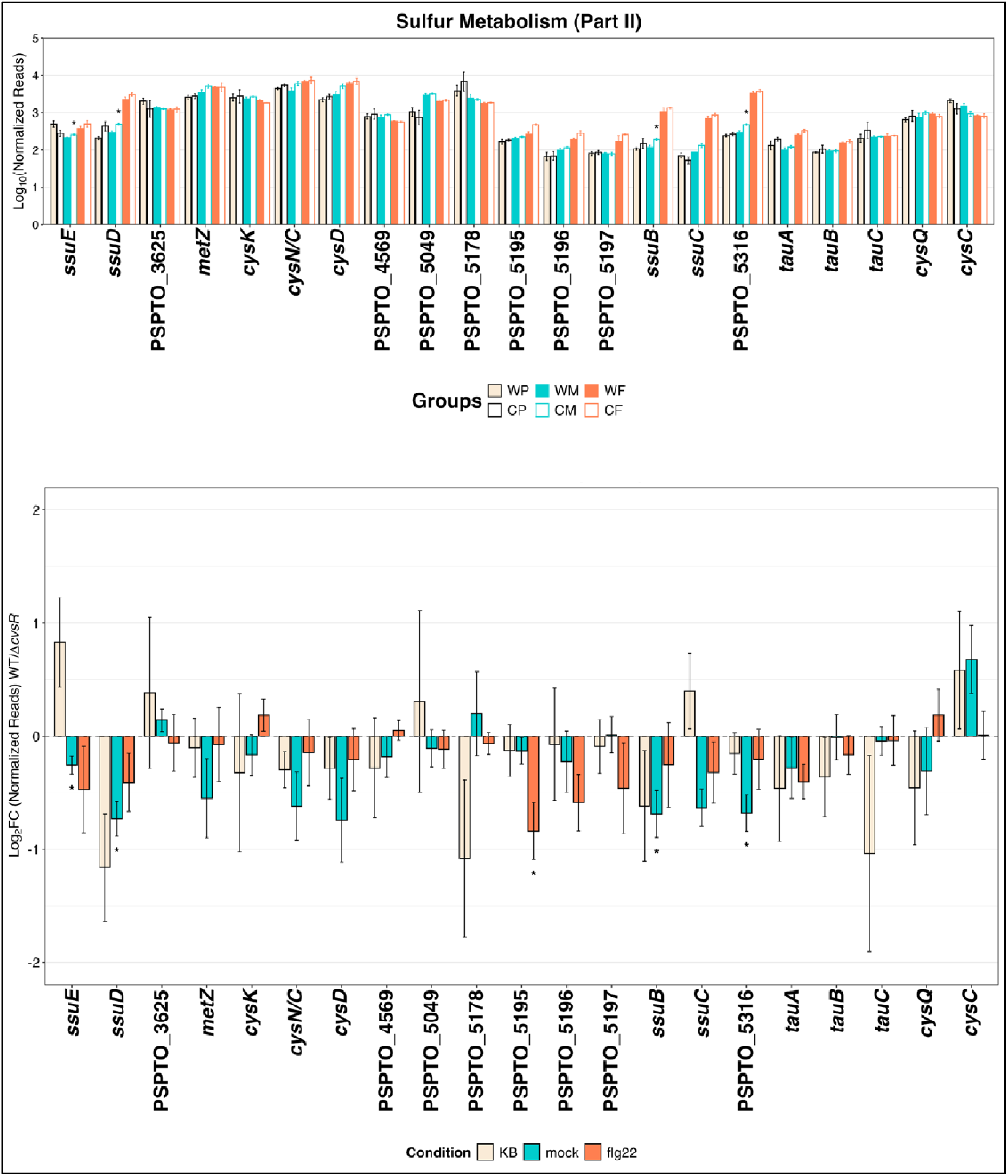
Expression of bacterial sulfur metabolism genes. (A) Normalized reads of gene listed in sulfate metabolism pathway of *P. syringae* pv. *tomato* DC3000 in each strain/condition: *cvsR* (C), wild type (W), flg22 treatment (F), and mock control (M). (B) log_2_ fold change in mean normalized gene counts of *cvsR* relative to wildtype of *in vitro* (KB), naïve tissue (mock), and PTI tissue (flg22). Asterisks indicate significant in changes for cvsR relative to wildtype (t.test, *P* > 0.05).

**Figure S4.**
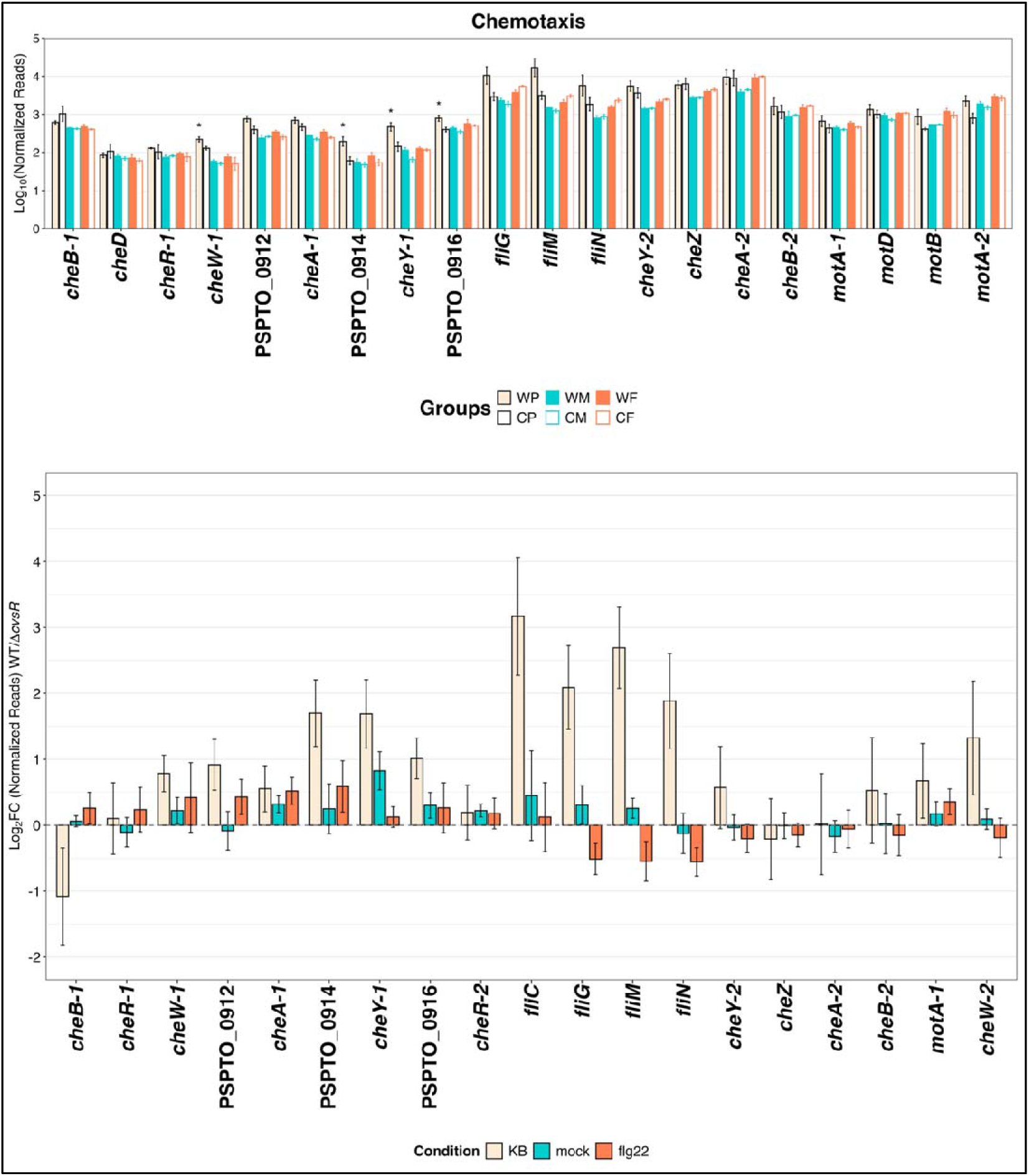
Expression of bacterial chemotaxis genes. (A) Normalized reads of gene listed in bacterial chemotaxis pathway of *P. syringae* pv. *tomato* DC3000 in each strain/condition: *cvsR* (C), wild type (W), flg22 treatment (F), and mock control (M). (B) log_2_ fold change in mean normalized gene counts of *cvsR* relative to wildtype of *in vitro* (KB), naïve tissue (mock), and PTI tissue (flg22). Asterisks indicate significant in changes for Δ*cvsR* relative to wildtype (t.test, *P* > 0.05).

**Figure S5.**
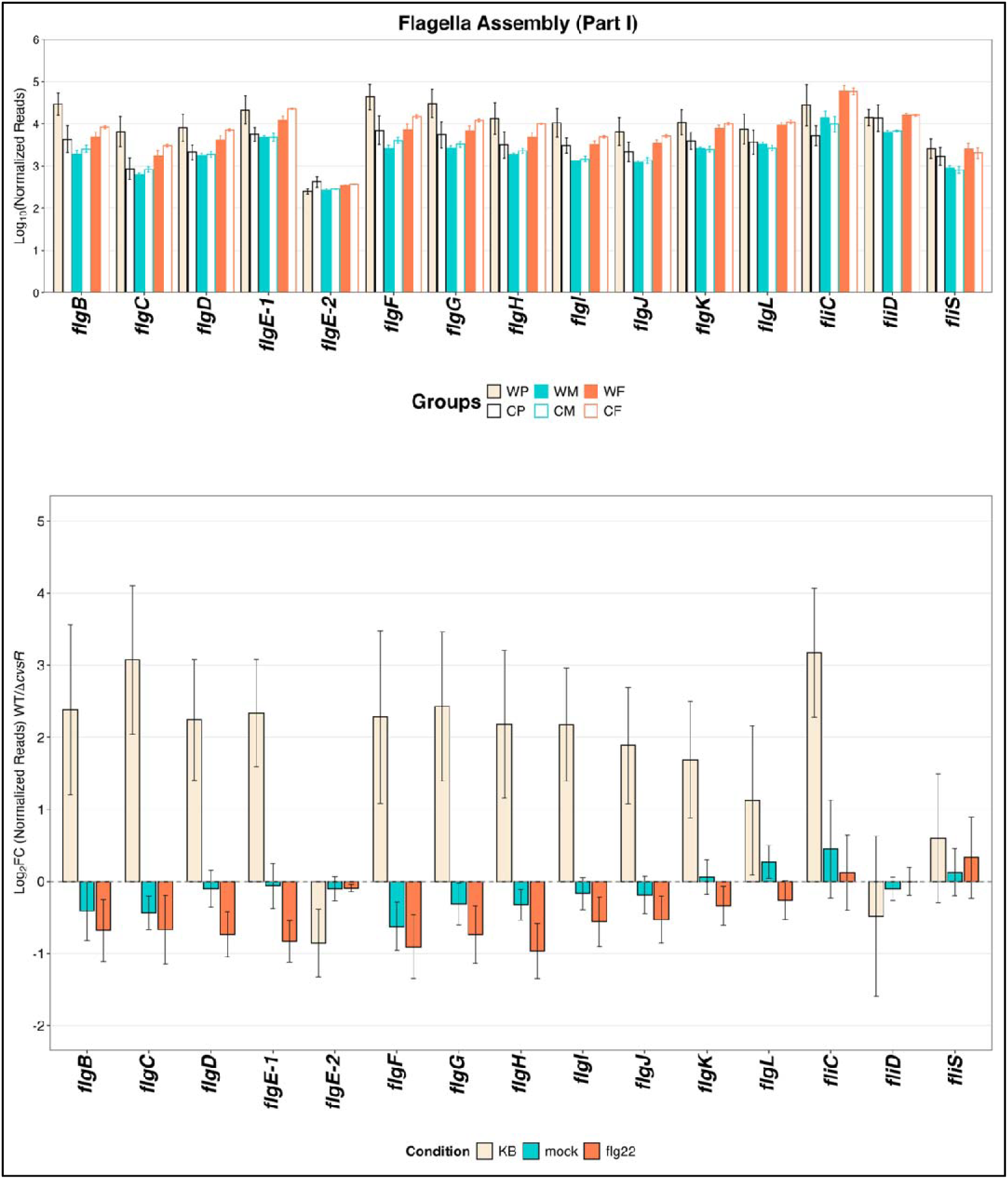
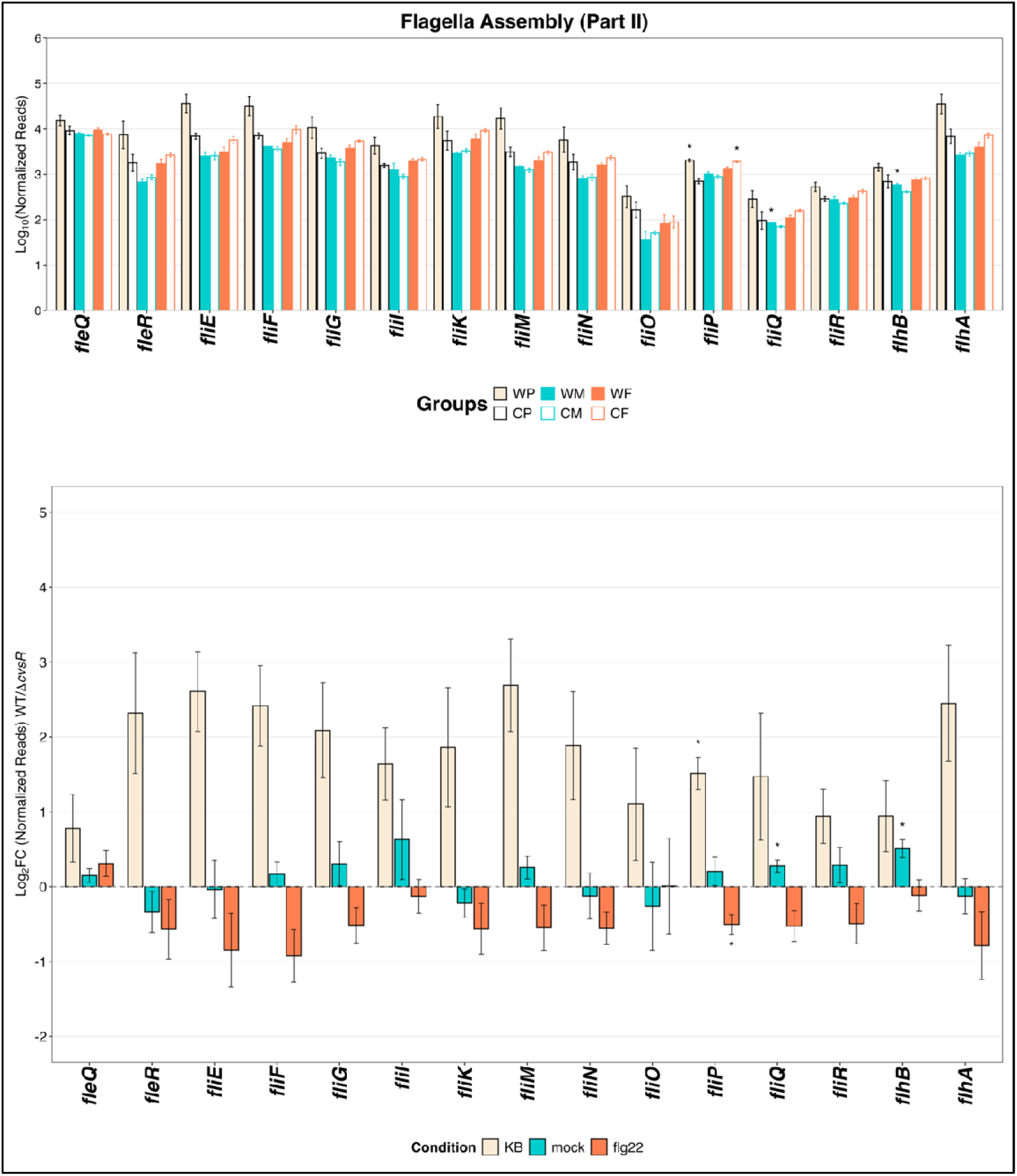
Expression of bacterial flagella assembly genes. (A) Normalized reads of gene listed in bacterial flagella assembly pathway of *P. syringae* pv. *tomato* DC3000 in each strain/condition: *cvsR* (C), wild type (W), flg22 treatment (F), and mock control (M). (B) log_2_ fold change in mean normalized gene counts of *cvsR* relative to wildtype of *in vitro* (KB), naïve tissue (mock), and PTI tissue (flg22). Asterisks indicate significant in changes for cvsR relative to wildtype (t.test, *P* > 0.05).

**Figure S6.**
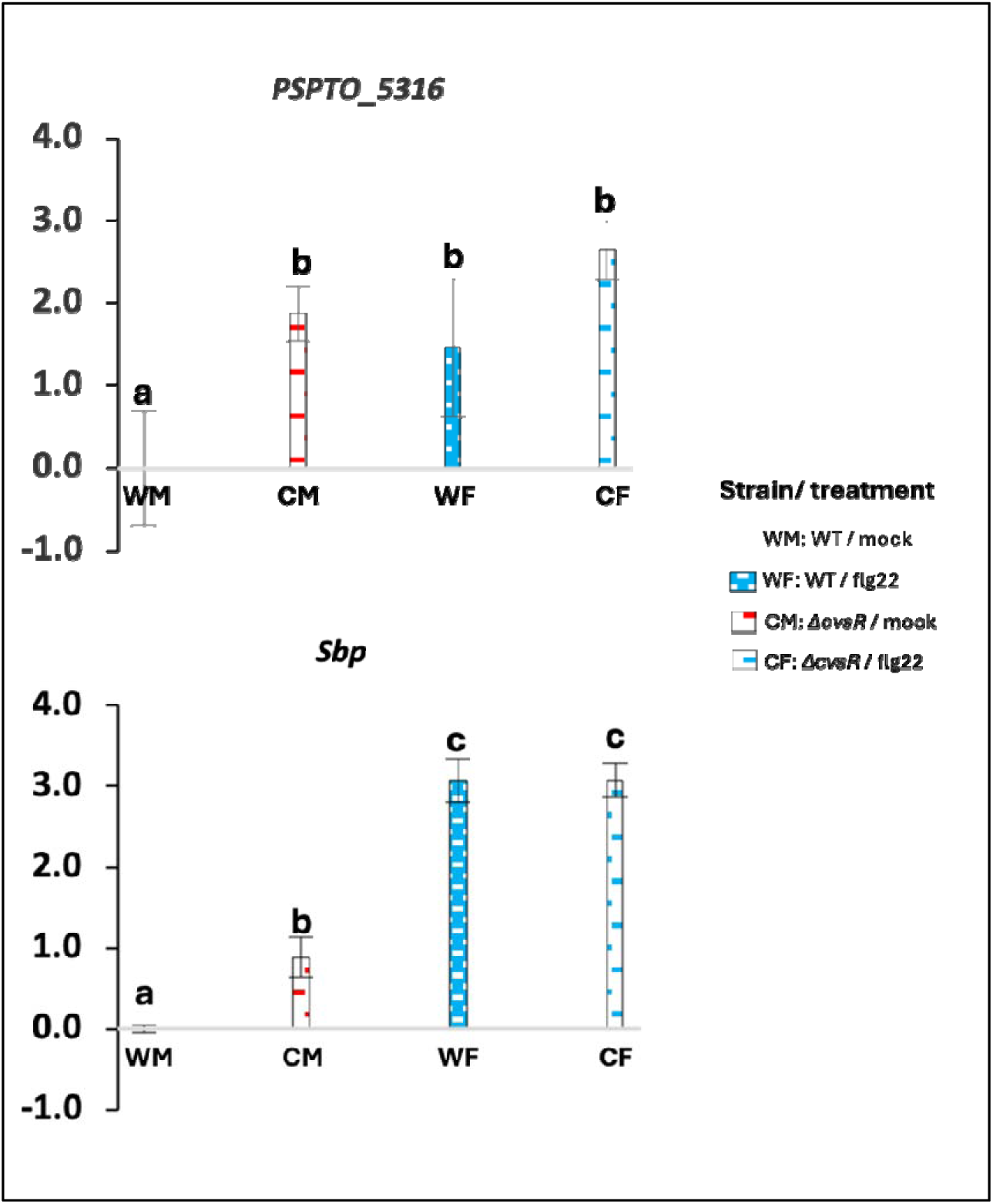
Reverse transcription-quantitative polymerase chain reaction (RT-qPCR) analysis of *Pto* DC3000 gene expression during early exposure to pattern-triggered immunity (PTI) relative to naïve host infection of *A. thaliana*. Expression of two sulfate metabolism–related genes (*sfbp* and *sbp*) in bacteria exposed to PTI relative to expression in bacteria during naïve host infection (PTI versus naïve), normalized against the expression of the reference gene *recA*. Bars represent mean log□normalized relative quantity (NRQ) and standard error calculated from independent biological replicate samples (n = 3). Different letters indicate statistically significant differences between conditions based on Student’s t-test (p < 0.05). Δ*cvsR* mutant (C), wild type (W), flg22 treatment (F), and mock control (M).

## References

Bolger, A. M., Lohse, M., & Usadel, B. (2014). Trimmomatic: A flexible trimmer for Illumina sequence data. Bioinformatics, 30(15), 2114–2120. 10.1093/bioinformatics/btu170

Buell, C. R., Joardar, V., Lindeberg, M., Selengut, J., Paulsen, I. T., Gwinn, M. L., Dodson, R. J., Deboy, R. T., Durkin, A. S., Kolonay, J. F., Madupu, R., Daugherty, S., Brinkac, L., Beanan, M. J., Haft, D. H., Nelson, W. C., Davidsen, T., Zafar, N., Zhou, L., … Collmer, A. (2003). The complete genome sequence of the Arabidopsis and tomato pathogen Pseudomonas syringae pv. Tomato DC3000. Proceedings of the National Academy of Sciences, 100(18), 10181–10186. 10.1073/pnas.1731982100

Chatterjee, A., Cui, Y., Yang, H., Collmer, A., Alfano, J. R., & Chatterjee, A. K. (2003). GacA, the Response Regulator of a Two-Component System, Acts as a Master Regulator in Pseudomonas syringae pv. Tomato DC3000 by Controlling Regulatory RNA, Transcriptional Activators, and Alternate Sigma Factors. Molecular Plant-Microbe Interactions®, 16(12), 1106–1117. 10.1094/MPMI.2003.16.12.1106

Chen, H.-C., Newton, C. J., Diaz, G., Zheng, Y., Kong, F., Yao, Y., Yang, L., & Kvitko, B. H. (2025). Proteomic snapshot of pattern triggered immunity in the Arabidopsis leaf apoplast. The Plant Journal, 123(6), e70498. 10.1111/tpj.70498

Chinchilla, D., Bauer, Z., Regenass, M., Boller, T., & Felix, G. (2006). The Arabidopsis Receptor Kinase FLS2 Binds flg22 and Determines the Specificity of Flagellin Perception. The Plant Cell, 18(2), 465–476. 10.1105/tpc.105.036574

Crabill, E., Joe, A., Block, A., van Rooyen, J. M., & Alfano, J. R. (2010). Plant immunity directly or indirectly restricts the injection of type III effectors by the Pseudomonas syringae type III secretion system. Plant Physiol, 154(1), 233–244. 10.1104/pp.110.159723

De Francesco, A., Lovelace, A. H., Shaw, D., Qiu, M., Wang, Y., Gurung, F., Ancona, V., Wang, C., Levy, A., Jiang, T., & Ma, W. (2022). Transcriptome Profiling of ‘Candidatus Liberibacter asiaticus’ in Citrus and Psyllids. Phytopathology®, 112(1), 116–130. 10.1094/PHYTO-08-21-0327-FI

Dellagi, A., Rigault, M., Segond, D., Roux, C., Kraepiel, Y., Cellier, F., Briat, J.-F., Gaymard, F., & Expert, D. (2005). Siderophore-mediated upregulation of Arabidopsis ferritin expression in response to Erwinia chrysanthemi infection. The Plant Journal, 43(2), 262–272. 10.1111/j.1365-313X.2005.02451.x

Deng, X., Liang, H., Chen, K., He, C., Lan, L., & Tang, X. (2014). Molecular mechanisms of two-component system RhpRS regulating type III secretion system in Pseudomonas syringae. Nucleic Acids Research, 42(18), 11472–11486. 10.1093/nar/gku865

Dora, S., Terrett, O. M., & Sánchez-Rodríguez, C. (2022). Plant–microbe interactions in the apoplast: Communication at the plant cell wall. The Plant Cell, 34(5), 1532–1550. 10.1093/plcell/koac040

Fatima, U., & Senthil-Kumar, M. (2021). Sweet revenge: AtSWEET12 in plant defense against bacterial pathogens by apoplastic sucrose limitation (p. 2021.10.04.463061). bioRxiv. 10.1101/2021.10.04.463061

Felix, G., & Boller, T. (2003). Molecular Sensing of Bacteria in Plants: THE HIGHLY CONSERVED RNA-BINDING MOTIF RNP-1 OF BACTERIAL COLD SHOCK PROTEINS IS RECOGNIZED AS AN ELICITOR SIGNAL IN TOBACCO *. Journal of Biological Chemistry, 278(8), 6201–6208. 10.1074/jbc.M209880200

Fishman, M. R., & Filiatrault, M. J. (2019). Prevention of Surface-Associated Calcium Phosphate by the Pseudomonas syringae Two-Component System CvsSR. Journal of Bacteriology, 201(7), 10.1128/jb.00584-18. 10.1128/jb.00584-18

Fishman, M. R., Zhang, J., Bronstein, P. A., Stodghill, P., & Filiatrault, M. J. (2018). Ca2+-Induced Two-Component System CvsSR Regulates the Type III Secretion System and the Extracytoplasmic Function Sigma Factor AlgU in Pseudomonas syringae pv. Tomato DC3000. Journal of Bacteriology, 200(5), 10.1128/jb.00538-17. 10.1128/jb.00538-17

Gómez-Gómez, L., Bauer, Z., & Boller, T. (2001). Both the Extracellular Leucine-Rich Repeat Domain and the Kinase Activity of FLS2 Are Required for Flagellin Binding and Signaling in Arabidopsis. The Plant Cell, 13(5), 1155–1164. https://pmc.ncbi.nlm.nih.gov/articles/PMC135565/

Guo, M., Tian, F., Wamboldt, Y., & Alfano, J. R. (2009). The majority of the type III effector inventory of Pseudomonas syringae pv. Tomato DC3000 can suppress plant immunity. Molecular Plant-Microbe Interactions: MPMI, 22(9), 1069–1080. 10.1094/MPMI-22-9-1069

Herlihy, J. H., Long, T. A., & McDowell, J. M. (2020). Iron homeostasis and plant immune responses: Recent insights and translational implications. Journal of Biological Chemistry, 295(39), 13444–13457. 10.1074/jbc.REV120.010856

Honda, T., Yu, S., Mai, D., Baumgart, L., Babnigg, G., & Yoshikuni, Y. (2025). CRAGE-RB-PI-seq enables transcriptional profiling of rhizobacteria during plant-root colonization (p. 2024.11.19.624340). bioRxiv. 10.1101/2024.11.19.624340

Jones, J. D. G., & Dangl, J. L. (2006). The plant immune system. Nature, 444(7117), 323–329. 10.1038/nature05286

Kim, N., Kim, J. J., Kim, I., Mannaa, M., Park, J., Kim, J., Lee, H.-H., Lee, S.-B., Park, D.-S., Sul, W. J., & Seo, Y.-S. (2020). Type VI secretion systems of plant-pathogenic Burkholderia glumae BGR1 play a functionally distinct role in interspecies interactions and virulence. Molecular Plant Pathology, 21(8), 1055–1069. 10.1111/mpp.12966

King, E. O., Ward, M. K., & Raney, D. E. (1954). Two simple media for the demonstration of pyocyanin and fluorescin. The Journal of Laboratory and Clinical Medicine, 44(2), 301–307.

Kunze, G., Zipfel, C., Robatzek, S., Niehaus, K., Boller, T., & Felix, G. (2004). The N Terminus of Bacterial Elongation Factor Tu Elicits Innate Immunity in Arabidopsis Plants. The Plant Cell, 16(12), 3496–3507. 10.1105/tpc.104.026765

Kvitko, B. H., Park, D. H., Velásquez, A. C., Wei, C.-F., Russell, A. B., Martin, G. B., Schneider, D. J., & Collmer, A. (2009). Deletions in the repertoire of Pseudomonas syringae pv. Tomato DC3000 type III secretion effector genes reveal functional overlap among effectors. PLoS Pathogens, 5(4), e1000388. 10.1371/journal.ppat.1000388

Langmead, B., & Salzberg, S. L. (2012). Fast gapped-read alignment with Bowtie 2. Nature Methods, 9(4), 357–359. 10.1038/nmeth.1923

Lavín, J. L., Kiil, K., Resano, O., Ussery, D. W., & Oguiza, J. A. (2007). Comparative genomic analysis of two-component regulatory proteins in Pseudomonas syringae. BMC Genomics, 8(1), 397. 10.1186/1471-2164-8-397

Lee, S. E., Gupta, R., Jayaramaiah, R. H., Lee, S. H., Wang, Y., Park, S. R., & Kim, S. T. (2017). Global Transcriptome Profiling of Xanthomonas oryzae pv. Oryzae under in planta Growth and in vitro Culture Conditions. Plant Pathol J, 33(5), 458–466. 10.5423/ppj.Oa.04.2017.0076

Lewis, L. A., Polanski, K., de Torres-Zabala, M., Jayaraman, S., Bowden, L., Moore, J., Penfold, C. A., Jenkins, D. J., Hill, C., Baxter, L., Kulasekaran, S., Truman, W., Littlejohn, G., Prusinska, J., Mead, A., Steinbrenner, J., Hickman, R., Rand, D., Wild, D. L., … Grant, M. (2015). Transcriptional Dynamics Driving MAMP-Triggered Immunity and Pathogen Effector-Mediated Immunosuppression in Arabidopsis Leaves Following Infection with Pseudomonas syringae pv tomato DC3000. The Plant Cell, 27(11), 3038–3064. 10.1105/tpc.15.00471

Liao, Z.-X., Ni, Z., Wei, X.-L., Chen, L., Li, J.-Y., Yu, Y.-H., Jiang, W., Jiang, B.-L., He, Y.-Q., & Huang, S. (2019). Dual RNA-seq of Xanthomonas oryzae pv. Oryzicola infecting rice reveals novel insights into bacterial-plant interaction. PLOS ONE, 14(4), e0215039. 10.1371/journal.pone.0215039

Love, M. I., Huber, W., & Anders, S. (2014). Moderated estimation of fold change and dispersion for RNA-seq data with DESeq2. Genome Biology, 15(12), 550. 10.1186/s13059-014-0550-8

Lovelace, A. H., Smith, A., & Kvitko, B. H. (2018). Pattern-Triggered Immunity Alters the Transcriptional Regulation of Virulence-Associated Genes and Induces the Sulfur Starvation Response in Pseudomonas syringae pv. Tomato DC3000. Molecular Plant-Microbe Interactions®, 31(7), 750–765. 10.1094/MPMI-01-18-0008-R

Luneau, J. S., Baudin, M., Quiroz Monnens, T., Carrère, S., Bouchez, O., Jardinaud, M.-F., Gris, C., François, J., Ray, J., Torralba, B., Arlat, M., Lewis, J. D., Lauber, E., Deutschbauer, A. M., Noël, L. D., & Boulanger, A. (2022). Genome-wide identification of fitness determinants in the Xanthomonas campestris bacterial pathogen during early stages of plant infection. New Phytologist, 236(1), 235–248. 10.1111/nph.18313

Luo, W., Friedman, M. S., Shedden, K., Hankenson, K. D., & Woolf, P. J. (2009). GAGE: Generally applicable gene set enrichment for pathway analysis. BMC Bioinformatics, 10(1), 161. 10.1186/1471-2105-10-161

Miao, P., Wang, H., Wang, W., Wang, Z., Ke, H., Cheng, H., Ni, J., Liang, J., Yao, Y.-F., Wang, J., Zhou, J.-M., & Lei, X. (2025). A widespread plant defense compound disarms bacterial type III injectisome assembly. Science, 387(6737), eads0377. 10.1126/science.ads0377

Ngou, B. P. M., Jones, J. D. G., & Ding, P. (2022). Plant immune networks. Trends in Plant Science, 27(3), 255–273. 10.1016/j.tplants.2021.08.012

Nobori, T., Velásquez, A. C., Wu, J., Kvitko, B. H., Kremer, J. M., Wang, Y., He, S. Y., & Tsuda, K. (2018). Transcriptome landscape of a bacterial pathogen under plant immunity. Proceedings of the National Academy of Sciences, 115(13), E3055–E3064. 10.1073/pnas.1800529115

Pertea, M., Kim, D., Pertea, G. M., Leek, J. T., & Salzberg, S. L. (2016). Transcript-level expression analysis of RNA-seq experiments with HISAT, StringTie and Ballgown. Nature Protocols, 11(9), 1650–1667. 10.1038/nprot.2016.095

Rogan, C. J., Pang, Y.-Y., Mathews, S. D., Turner, S. E., Weisberg, A. J., Lehmann, S., Rentsch, D., & Anderson, J. C. (2024). Transporter-mediated depletion of extracellular proline directly contributes to plant pattern-triggered immunity against a bacterial pathogen. Nature Communications, 15(1), 7048. 10.1038/s41467-024-51244-6

Shao, X., Tan, M., Xie, Y., Yao, C., Wang, T., Huang, H., Zhang, Y., Ding, Y., Liu, J., Han, L., Hua, C., Wang, X., & Deng, X. (2021). Integrated regulatory network in Pseudomonas syringae reveals dynamics of virulence. Cell Reports, 34(13), 108920. 10.1016/j.celrep.2021.108920

Smith, A., Lovelace, A. H., & Kvitko, B. H. (2018). Validation of RT-qPCR Approaches to Monitor Pseudomonas syringae Gene Expression During Infection and Exposure to Pattern-Triggered Immunity. Molecular Plant-Microbe Interactions®, 31(4), 410–419. 10.1094/MPMI-11-17-0270-TA

Sreedharan, A., Penaloza-Vazquez, A., Kunkel, B. N., & Bender, C. L. (2006). CorR Regulates Multiple Components of Virulence in Pseudomonas syringae pv. Tomato DC3000. Molecular Plant-Microbe Interactions®, 19(7), 768–779. 10.1094/MPMI-19-0768

Wang, H., Smith, A., Lovelace, A., & Kvitko, B. H. (2022). In planta transcriptomics reveals conflicts between pattern-triggered immunity and the AlgU sigma factor regulon. PLOS ONE, 17(9), e0274009. 10.1371/journal.pone.0274009

Wang, N., Han, N., Tian, R., Chen, J., Gao, X., Wu, Z., Liu, Y., & Huang, L. (2021). Role of the Type VI Secretion System in the Pathogenicity of Pseudomonas syringae pv. Actinidiae, the Causative Agent of Kiwifruit Bacterial Canker. Frontiers in Microbiology, 12. 10.3389/fmicb.2021.627785

Whalen, M. C., Innes, R. W., Bent, A. F., & Staskawicz, B. J. (1991). Identification of Pseudomonas syringae pathogens of Arabidopsis and a bacterial locus determining avirulence on both Arabidopsis and soybean. The Plant Cell, 3(1), 49–59. 10.1105/tpc.3.1.49

Xin, X.-F., Kvitko, B., & He, S. Y. (2018). Pseudomonas syringae: What it takes to be a pathogen. Nature Reviews Microbiology, 16(5), 316–328. 10.1038/nrmicro.2018.17

Yamada, K., & Mine, A. (2024). Sugar coordinates plant defense signaling. Science Advances, 10(4), eadk4131. 10.1126/sciadv.adk4131

Yang, S., Lovelace, A. H., Yuan, Y., Nie, H., Chen, W., Gao, Y., Bo, W., Nagel, D. H., Pang, X., & Ma, W. (2025). A witches’ broom phytoplasma effector induces stunting by stabilizing a bHLH transcription factor in Ziziphus jujuba plants. New Phytologist, 247(1), 249–264. 10.1111/nph.70172

Yang, Z., Wang, H., Keebler, R., Lovelace, A., Chen, H.-C., Kvitko, B., & Swingle, B. (2024). Environmental alkalization suppresses deployment of virulence strategies in Pseudomonas syringae pv. Tomato DC3000. Journal of Bacteriology, 206(11), e00086–24. 10.1128/jb.00086-24

Yu, K., Liu, Y., Tichelaar, R., Savant, N., Lagendijk, E., Kuijk, S. J. L. van, Stringlis, I. A., Dijken, A. J. H. van, Pieterse, C. M. J., Bakker, P. A. H. M., Haney, C. H., & Berendsen, R. L. (2019). Rhizosphere-Associated Pseudomonas Suppress Local Root Immune Responses by Gluconic Acid-Mediated Lowering of Environmental pH. Current Biology, 29(22), 3913–3920.e4. 10.1016/j.cub.2019.09.015

Yuan, M., Ngou, B. P. M., Ding, P., & Xin, X.-F. (2021). PTI-ETI crosstalk: An integrative view of plant immunity. Current Opinion in Plant Biology, 62, 102030. 10.1016/j.pbi.2021.102030

Zhou, J.-M., & Zhang, Y. (2020). Plant Immunity: Danger Perception and Signaling. Cell, 181(5), 978–989. 10.1016/j.cell.2020.04.028

Zipfel, C., Robatzek, S., Navarro, L., Oakeley, E. J., Jones, J. D. G., Felix, G., & Boller, T. (2004). Bacterial disease resistance in Arabidopsis through flagellin perception. Nature, 428(6984), 764–767. 10.1038/nature02485

